# An SMR cell-cycle inhibitor inducible by a carotenoid metabolite resets root development and drought tolerance in Arabidopsis

**DOI:** 10.1101/2023.05.12.540478

**Authors:** Jeanne Braat, Meryl Jaonina, Pascale David, Maïté Leschevin, Bertrand Légeret, Stefano D’Alessandro, Frédéric Beisson, Michel Havaux

**Author notes:** Correspondence to Michel Havaux.

## Abstract

New regulatory functions in plant development and environmental stress responses have recently emerged for a number of apocarotenoids produced by enzymatic or non-enzymatic oxidation of carotenoids. β-cyclocitric acid (β-CCA) is one such compound derived from β-carotene which triggers defense mechanisms leading to a marked enhancement of plant tolerance to drought stress. We show here that this response is associated with an inhibition of root growth affecting both root cell elongation and division. Remarkably, β-CCA selectively induced cell cycle inhibitors of the SIAMESE-RELATED (SMR) family, especially SMR5, in root tip cells. Overexpression of the *SMR5* gene in Arabidopsis induced molecular and physiological changes that mimicked in large part the effects of β-CCA. In particular, the *SMR5* overexpressors exhibited an inhibition of root development and a marked increase in drought tolerance which is not related to stomatal closure. *SMR5* up-regulation induced changes in gene expression that strongly overlapped with the β-CCA-induced transcriptomic changes. Both β-CCA and SMR5 led to a down-regulation of many cell cycle activators (cyclins, cyclin-dependent kinases) and a concomitant up-regulation of genes related to water deprivation, cellular detoxification and biosynthesis of lipid biopolymers such as suberin and lignin. This was correlated with an accumulation of suberin lipid polyesters in the roots and a decrease in non-stomatal leaf transpiration. Taken together, our results identify the β-CCA-and drought-inducible *SMR5* gene as a key component of a stress signaling pathway that reorients root metabolism from growth to multiple defense mechanisms leading to drought tolerance.

## Introduction

Carotenoids are a large group of compounds constituted by eight isoprene units, the vast majority of which are derived from the linear tetraterpene phytoene (DellaPenna and Pogson 2006, Ruiz-Sola and Rodriguez-Concepcion 2012, Nisar et al. 2015, Sun et al. 2022). Modifications of this linear backbone by desaturases, cyclases, hydroxylases, ketolases and other enzymes give rise to a wide diversity of compounds. The presence of conjugated double bonds in the carotenoid skeleton confers pigment properties to this family of molecules, allowing them to play a variety of physiological functions. Plant carotenoids are mainly known as accessory light-harvesting pigments in photosynthesis, antioxidants, lipid membrane stabilizers and attractants for pollinators and seed dispersers. Carotenoids are also at the origin of a number of bioactive molecules, including phytohormones such as strigolactones and abscisic acid, by enzymatic or non-enzymatic oxidative cleavage (Nambara and Marion-Poll 2005, Bouwmeester et al. 2019). New regulatory functions have recently emerged for some of those cleavage products (apocarotenoids) which relate to plant development and response to environmental stresses (Havaux 2014, Moreno et al. 2021, Sierra et al. 2022). The volatile β-cyclocitral (β-CC), generated by the oxidation of β-carotene, is one such bioactive compound, which is partly converted *in planta* into the water-soluble β-cyclocitric acid (β-CCA) (D’Alessandro and Havaux 2019, Havaux 2020). β-CC and its oxidized form β-CCA function in plants as molecular signals causing metabolic changes and triggering changes in nuclear gene expression which lead to an increased tolerance to several biotic and abiotic stresses (Ramel et al. 2012, D’Alessandro et al. 2019, Mitra et al. 2021). In particular, increasing the internal concentration of β-CCA in plants by exogenous applications was shown to bring about a marked enhancement of drought tolerance (D’Alessandro et al. 2019).

Transcriptomic analyses of plant leaves exposed to volatile β-CC have shown that this apocarotenoid induces cellular defense and detoxification mechanisms (Ramel et al. 2012, Deshpande et al. 2021). More precisely, β-CC triggers the so-called xenobiotic detoxification pathway controlled by TGAII transcription factors interacting with the SCL14 transcription regulator (D’Alessandro et al. 2018). The TGAII/SCL14 complex governs the expression of a variety of detoxification enzymes which target toxic reactive carbonyls such as those produced by lipid peroxidation (Fode et al. 2008, Mueller et al. 2008). Enhancement of the detoxification capacities by β-CC(A) is likely to participate in the enhancement of plant tolerance to stressful conditions, such as drought stress, that produce reactive oxygen species and lead to lipid peroxidation (D’Alessandro et al. 2019, Rac et al. 2022).

β-CC was also reported to promote root growth in Arabidopsis, rice and tomato (Dickinson et al. 2019). This effect was maintained under salt stress, suggesting that regulation of root stem cell behavior by β-CC could increase plant vigor under stressful conditions. This hypothesis prompted us to examine if β-CCA-induced drought stress tolerance is associated with changes in root growth. The results shown in this study indicate that, contrary to what was previously reported for β-CC, β-CCA affects root development by causing a marked inhibition of primary and secondary root growth. The detailed analysis of this phenomenon allowed us to identify a gene, inducible by β-CCA and water stress, whose expression largely mimics the effects of β-CCA on transcriptomic responses, root growth and drought tolerance. The present study describes this new key component of the β-CCA signaling pathway.

## Results

### The induction of drought tolerance by β-CCA is associated with root growth inhibition

Arabidopsis plants grown on soil were watered with 1.5 mM β-CCA (or with water for the controls) before stopping watering for 10 d. As expected (D’Alessandro et al. 2019), plants pretreated with β-CCA were much more tolerant to water deprivation than control plants (Fig. 1a). β-CCA-treated plants remained fully turgid while control plants showed clear signs of leaf dehydration after 10 d of water deprivation. Rather surprisingly, the protective effect of β-CCA was associated with a marked reduction of root length (Fig 1b,c). Root growth inhibition was also observed when well-watered plants were treated with β-CCA and let to grow for 10 d in the absence of any water stress (Fig. 1d,e). Thus, reduction of root development appears to be a direct effect of β-CCA and does not necessarily require an interaction with drought stress. Fig. 1f shows that the reduction of root biomass was observed when plants were watered with β-CCA solutions at concentrations above 250 µM.

**Fig. 1.**
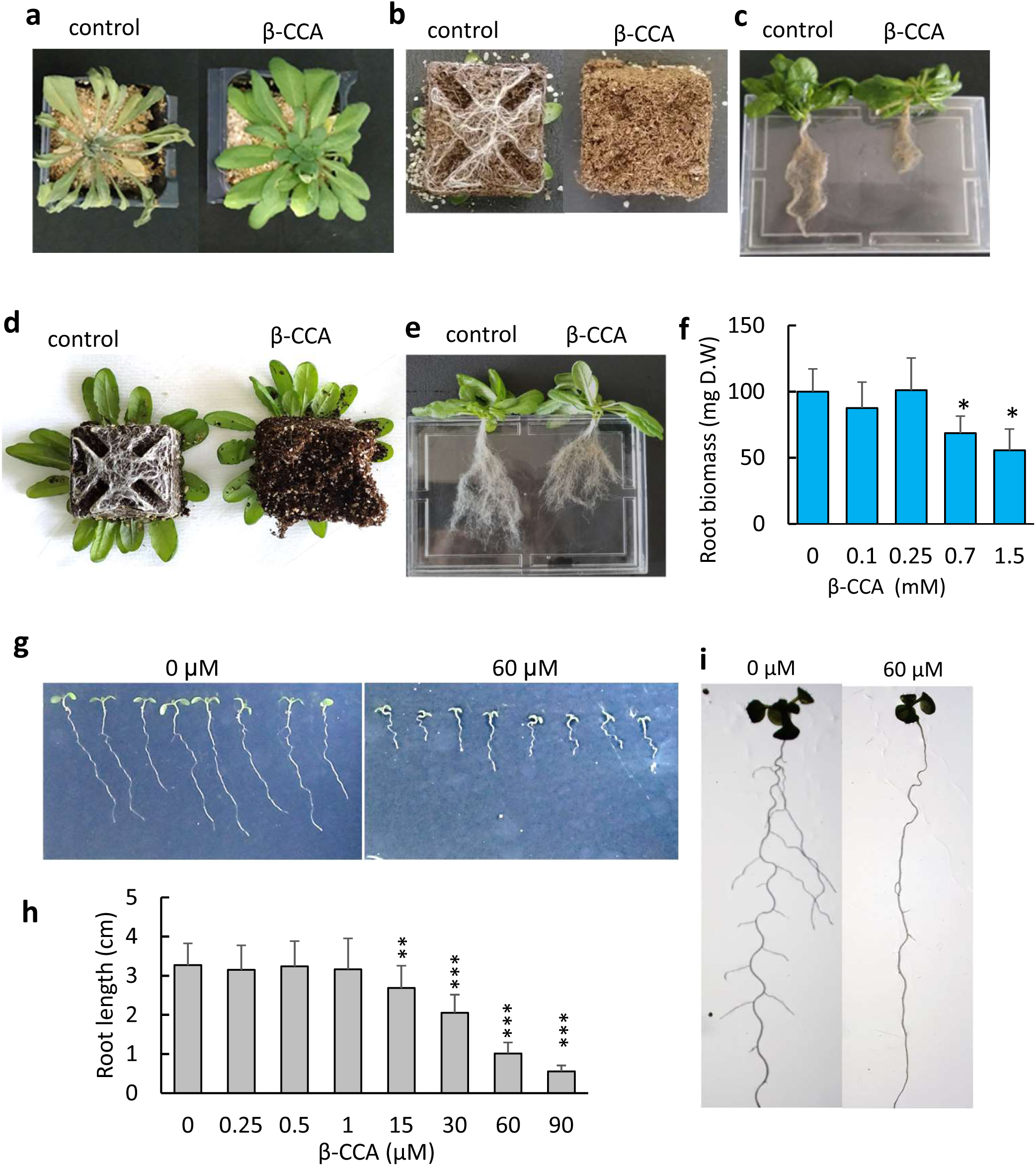
β-CCA inhibits root growth of Arabidopsis plants. A) Picture of Arabidopsis plants exposed to water stress induced by withdrawing watering for 10 d. β-CCA-pretreated plants were watered with 25 mL of a 1.5 mM β-CCA solution prior to water stress. Control plants received pure water instead of β-CCA. B) Picture of the roots at the bottom of the pot after 10 d of water stress, showing the decreased density of root hair with β-CCA treatment. C) Picture of the root system. D)-E) Picture of the roots of plants pretreated with 25 mL of 1.5 mM β-CCA or with 25 mL of water (control) and then let to growth with normal irrigation (no water stress). F) Root dry weight of plants treated with 25 ml of different solutions of β-CCA (0, 0.1, 0.25, 0.7, 1.5 mM β-CCA) in the absence of water stress. Data are mean values of 3 to 4 measurements + SD. *, different from 0 mM with P<0.05 (Student’s t-test). G) Picture of the seedlings grown *in vitro* on Agar with 0 or 60 µM β-CCA. H) Root length as a function of the β-CCA concentration in the invitro growth medium. Data are mean values of 15 measurements + SD. ** and ***, different from 0 µM with P < 0.01 and 0.001, respectively (Student’s t-test). I) Both the primary root and lateral roots of *in vitro*-grown seedlings were inhibited by β-CCA.

The effect of β-CCA on Arabidopsis root growth was further studied in seedlings grown on solid growth medium in Petri dishes. The results shown in Fig. 1g confirm the inhibitory action of the apocarotenoid on primary root length, which was observed at concentrations higher than 1 µM (Fig. 1h). No stimulatory effect was found in the nM range, contrary to what was previously reported for β-CC (Dickinson et al. 2019). Formation of lateral roots was also inhibited by β-CCA (Fig. 1i).

### Effects of β-CCA on root cell elongation and division

Root length is determined by cell proliferation in the meristem and cell expansion during differentiation. We measured both phenomena by microscopic analyses. First, we measured the effect of β-CCA on cell elongation in Arabidopsis roots colored with Ruthenium Red. Cell elongation was substantially reduced by β-CCA (Fig. 2a), and the dependence of this inhibition on the β-CCA concentration (Fig. 2b) was quite similar to the plot of root length vs. β-CCA concentration shown in Fig. 1h. We also measured the hypocotyl length of Arabidopsis seedlings grown in the dark, a phenomenon specifically due to cell elongation (Stuart et al. 1977, Gendreau et al. 1997). The results (Fig. 2c) show that β-CCA leads to shorter hypocotyls, confirming that the apocarotenoid is an inhibitor of cell expansion.

**Fig. 2.**
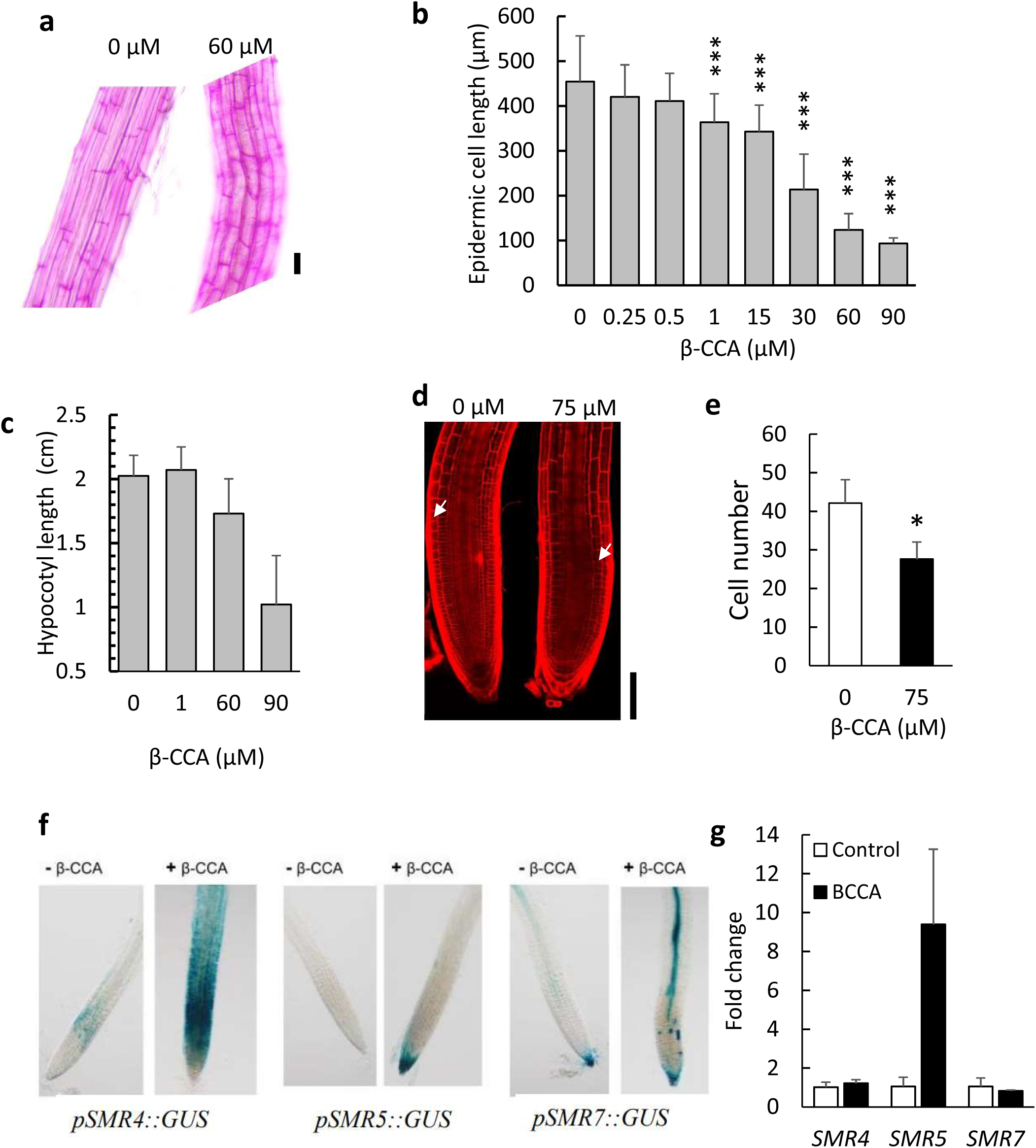
Both cell elongation and cell division in primary roots are perturbed by β-CCA. A) Picture of root epidermal cells in the elongation zone colored with Ruthenium Red. Seedlings were grown for 6 d on solid Agar medium with 0 or 60 µM β-CCA. B) Quantification of root cell length as a function of the β-CCA concentration. Data are mean values of 20 measurements + SD. ***, different from 0 µM with P<0.001 (Student’s t-test). C) Effect of β-CCA on Arabidopsis hypocotyl elongation in the dark. D) Picture of the meristem zone in Arabidopsis seedlings visualized by the fluorescence of Propridium Iodide. Arrows indicate the boudary between cell division and elongation zones. E) Meristem cortex cell number in the division zone of roots of Arabidopsis seedlings treated with 0 or 75 µM β-CCA. Cells were counted from the quiescent center to the boundary of the cell division and elongation zones. Values are means of 6 measurements + SD. *, p<0.05 in two-tailed Student’s test. Scale bar in panels a and c = 100 µm. F) GUS coloration of root tips of the transcriptional GUS reporter lines *pSMR4::GUS, pSMR5::GUS, pSMR7::GUS*. Seedlings were exposed to 0 or 75 µM β-CCA in the growth medium for 6 d. G) qRT-PCR analysis of the relative transcript levels of *SMR4, SMR5* and *SMR7*. Data are normalized to the housekeeping gene *UPL7* and are expressed as average fold changes + SD (3 replicates) compared to the control levels set to 1.

We also used confocal microscopy and fluorescence staining of cell walls with Propidium Iodide to investigate the effects of β-CCA on cell division in the root meristem (Fig. 2d). β-CCA markedly decreased the size of the root apical meristem. The average number of meristematic cortex cells in the apical meristem of β-CCA-treated seedlings was around 65% of control seedlings (Fig. 2e). This indicates a reduction of the cell division rate as a response to β-CCA.

Cell proliferation occurs as a result of periodic activation of cyclin-dependent kinases (CDK) by different cyclins (CYC), ensuring the transition from one phase of the mitotic cycle to another (De Veylder et al. 2003, Francis 2007, Tank and Thaker 2011). To characterize further the impact of β-CCA on root cell division, we checked the effect of β-CCA on the promoter activity of a panel of mitotic marker genes fused to a gene encoding the GUS reporter. These lines are markers of the G1 and S phases of the interphase (CYCA3 and CYCD) and of the second gap phase G2 and the M mitotic phase (CYCA2 and CYCB) (Bulankova et al 2013, Collins et al 2015, Forzani et al. 2014, Sozzani et al 2010). The color patterns of those GUS lines for cyclins are shown in Extended Data Fig. 1. No significant change was found for most cyclins (CYCA3;1, CYCD3;3, CYCB1;2, CYCA2;3, CYCA3;2, CCSS2A1) in response to β-CCA. An increase, rather than a decrease, in GUS coloration with β-CCA treatment was observed for CYCD6;1 and CYCB1;1. The effect of β-CCA on the expression of other cyclins and of CDKs will be shown below with the transcriptomic data.

CYC-CDK activity is regulated by several mechanisms, including transcriptional regulation, proteolysis and interactions with CDK inhibitors (Kumar and Larkin 2017). CDK inhibitors are crucial for plant development, particularly during the transition from the mitotic cell cycle to endoreplication (De Veylder et al. 2011, Inagaki and Umeda 2011). We examined the expression of the plant-specific CDK inhibitors SIAMESE (SIM) and SIAMESE-RELATED (SMR), which can interact with and inhibit all CDKs (Churchman et al. 2006, De Veylder et al. 2011, Kumar et al. 2015,Van Leene et al. 2010), using promoter-driven GUS reporter lines (Barrada et al. 2019). The expression of SIM did not respond to β-CCA (Extended Data Fig. 1). In contrast, SMR4, SMR5 and SMR7 GUS reporter lines showed the most remarkable responses (Fig. 2f). Their expression patterns were markedly enhanced upon β-CCA treatment. Intense and extensive staining encompassed both the division zone (SMR4, SMR5, SMR7) and the differentiation zone (SMR4 and SMR7). Although the SMR protein family is rather large (Kumar and Larkin 2017), SMR5, and to a lesser extent SMR4 and SMR7, have been previously shown to be the most responsive SMRs to environmental factors (Yi et al. 2014, Dubois et al. 2018).

We also analyzed the expression of the three *SMR* genes by qRT-PCR. *SMR5* transcripts (AT1G07500) accumulated when Arabidopsis was treated with β-CCA whereas no significant change was observed for the *SMR4* (AT5G02220) and *SMR7* (AT3G27630) transcripts (Fig. 2g). This differential response of the three *SMRs* was also found in the RNAseq analysis (see below). It is possible that the combination of low expression and very localized expression pattern of *SMR4* and *SMR7* makes them difficult to be monitored in the root samples harvested for the qRT-PCR analyses. The stability of the mRNA of the three *SMRs* may also be different.

The gene expression data of Fig. 2 shows that β-CCA induces the *SMR5* gene. Considering the known function of SMRs in cell division, this is potentially an important component of the root growth inhibition by β-CCA and possibly also of the enhancement of plant drought tolerance.

### Role of SMR4 and SMR5 in root growth

To examine the possible role of SMR expression in root growth and drought tolerance, we investigated the responses of SMR-deficient and -overexpressing Arabidopsis lines to water stress and β-CCA.

Suppression of *SMR4*, *SMR5* and *SMR7* in a triple mutant (*smr4 smr5 smr7*), previously described in Hendrix et al. (2020), had virtually no effect on shoot and root growth (Extended Data Fig. 2). Neither the shoot morphology and size (Extended Data Fig. 2a,b), nor the root length and biomass (Extended Data Fig. 2c-f) were affected by the lack of SMR4, SMR5 and SMR7. This was observed under different growth conditions: on soil, on sand and *in vitro* on solid Agar medium. Also, the response to β-CCA was not impaired in the *smr4 smr5 smr7* triple mutant: β-CCA inhibited root growth (Extended Data Fig. 2f) and enhanced drought tolerance (Extended Data Fig. 2g).

The SIM/SMR family is a large family, composed of at least 17 members in Arabidopsis (Kumar and Larkin 2017), and therefore we cannot exclude a compensation mechanism in the triple SMR mutant involving functional substitution by several members of the SIM/SMR family. There are also 7 genes encoding KRP proteins which constitute another class of CDK inhibitors (Kumar and Larkin 2017). To overcome the high redundancies in this regulation, we adopted an overexpression approach. Two lines overexpressing either *SMR4* or *SMR5* (designated as *OE:SMR4* or *OE:SMR5*), previously described by Yi et al. (2014), were available for this study. Both lines exhibited high levels of *SMR* expression compared to the control level (Fig. 3f).

**Fig. 3.**
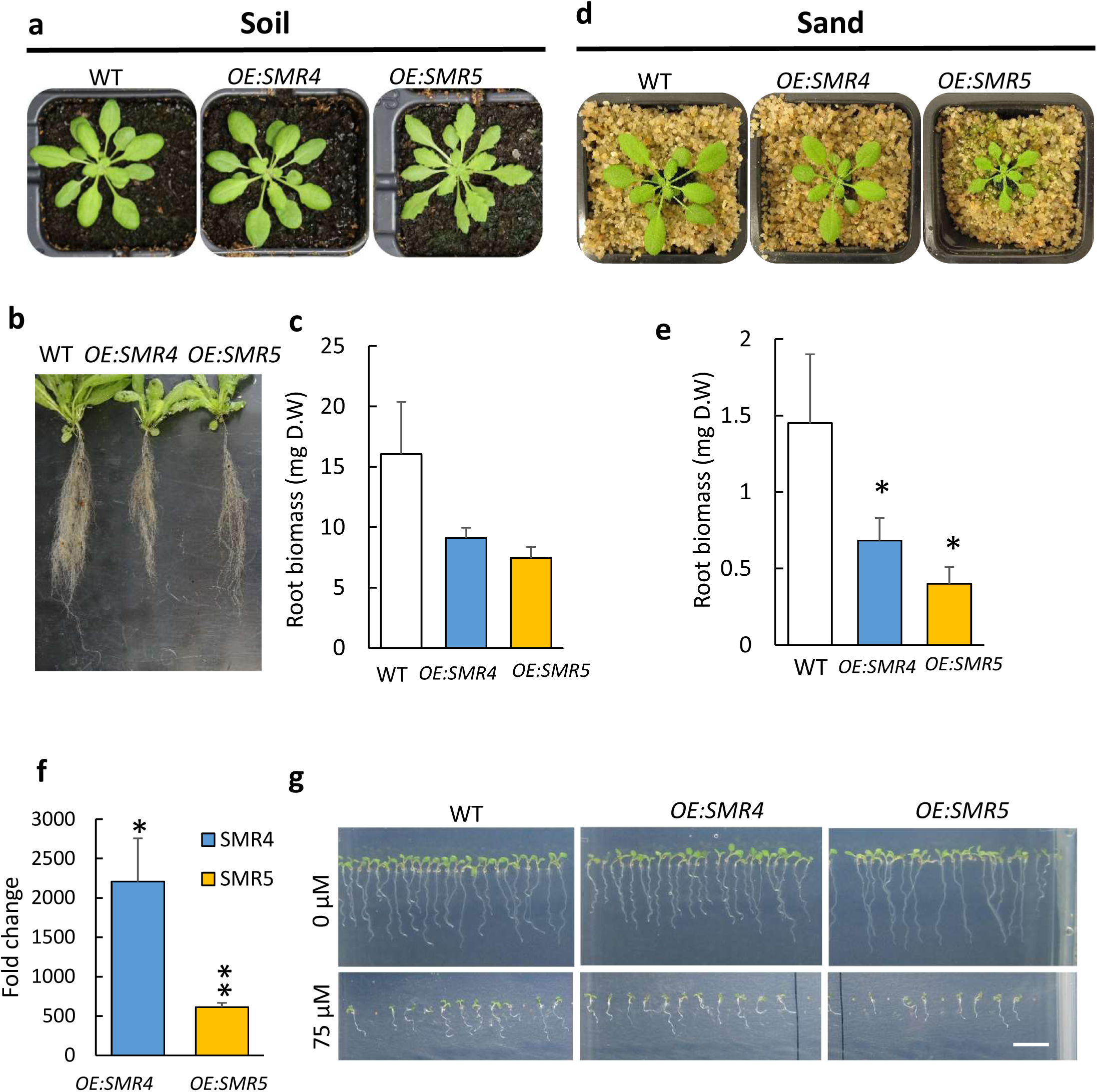
Growth of plants overexpressing SMR5 or SMR4. A) Expression levels of *SMR4* and *SMR5* in leaves of the corresponding overexpressing lines compared to WT elevls set to 1. B and E) Picture of the plants (WT, *OE:SMR5* and *OE:SMR4*) aged 5 weeks or 4 weeks grown on soil or on sand, respectively. c) Root system of plants grown on soil for 6 weeks. D and F) Root dry weight of plants grown on soil or on sand (3 to 7 replicates + SD). g) *In vitro* growth of the plants on Agar with or without 75 µM β-CCA. Scale bar, 1 cm. * and **, different from WT at P<0.05 and 0.01 respectively (Student’s t test).

When plants were grown on soil in our growth conditions, shoot development of the *SMR* overexpressors did not strongly differ from that of WT (Fig. 3a). There was no reduction of the fresh weight of shoot biomass of the *OE:SMRs* compared to WT (Extended Data Fig. 3). We observed a change in the shoot morphology of the *OE:SMR5* overexpressor which exhibited serrated leaves, as also reported previously by Yi et al. (2014). This seems to be a characteristics of plants exhibiting high SMR levels; it was also observed previously in a strong *SMR1* overexpressor containing elevated amounts of the SMR1 protein (Dubois et al. 2018).

Root development was significantly affected in *OE:SMR* plants, with a decrease in root biomass (dry weight), particularly in *OE:SMR5* relative to WT (Fig. 3b,c). Root growth was slightly less inhibited in *OE:SMR4* compared to *OE:SMR5* (Fig. 3c). Contrary to root biomass, the length of the primary root was not reduced in the *OE:SMR* lines (Fig. 3b), indicating only partially overlapping changes with those induced by β-CCA (Fig. 1). When plants were grown on sand, the growth phenotype of *OE:SMR4* and *OE:SMR5* was more marked, with root and shoot sizes being reduced in both overexpressors compared to WT (Fig. 3d,e). Thus, the impact of *SMR4* and *SMR5* overexpression appeared to be dependent on the growth conditions. This is confirmed when seedlings were grown *in vitro* on solid medium (Fig. 3g). Primary root growth was slightly inhibited in *OE:SMR4*, not in *OE:SMR5*. The environmental control of the overexpressor phenotype could be due to post-translational regulations that may affect the protein levels, as previously shown for *SMR1* (Dubois et al. 2018).

### Drought tolerance associated with SMR4 or SMR5 overexpression

WT and the two *SMR* overexpressors were exposed to drought stress imposed by withholding watering. Strikingly, *OE:SMR5* and *OE:SMR4* were much more tolerant to water stress than WT (Fig. 4a). After 10 d of water deprivation, WT plants were dehydrated, with the RWC dropping to ca. 40%, while both overexpressors remained turgescent, with a RWC above 90% (Fig. 4b). Drought tolerance of *OE:SMR4* was intermediate between WT and *OE:SMR5*: RWC decreased to the WT level at day 12. Strikingly, a prolonged water stress of 14 d was necessary to observe stress consequences in *OE:SMR5* plants, with RWC decreasing to around 50%.

To confirm the involvement of SMR4 and SMR5 in drought tolerance, we generated additional *SMR4-* and *SMR5*-overexpressing transgenic lines. Extended Data Fig. 4 presents a selection of independent transgenic lines with different levels of *SMR4* or *SMR5* overexpression (Extended Data Fig. 4b) after 12 d of water deprivation (Extended Data Fig. 4a). All the lines were more tolerant to water stress than WT, remaining turgescent and keeping high RWC values under very harsh conditions (Extended Data Fig. 4e). Root hair density was also reduced in all *OE:SMR* lines compared to WT (Extended Data Fig. 4c and d). It is to be noted that the majority of transgenic plants had serrated leaves.

**Fig. 4.**
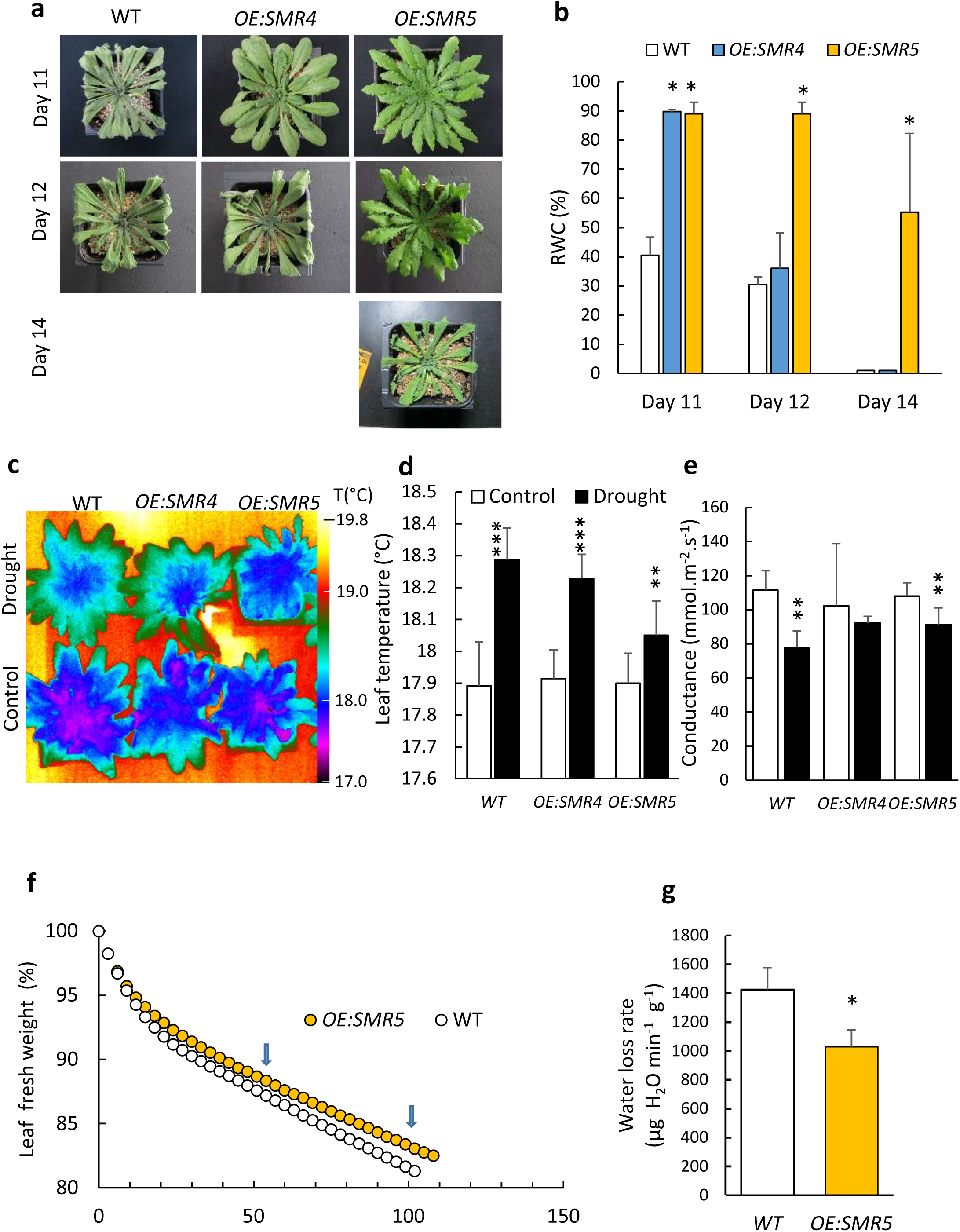
*SMR4-or SMR5*-overexpressing plants are tolerant to drought stress. a) Picture of the plants (WT, OE:SMR5 and OE:SMR4) water-stressed for 10, 12 and 14 days. Water stress was imposed by stopping watering. Absence of picture means that the plants died. B) Leaf RWC (mean values of 3 separate experiments + SD). *, different from WT at P<0.05 (Student’s t test). C) IR images of leaf temperatures in control plants and plants exposed to drought stress for 6 d. D) Average temperature of mature leaves. Data are mean values of 4 different plants + SD. E) Porometric measurements of stomatal conductance (6 replicates + SD). F-G) Water losses by excised plants in the dark. Data of panel e are mean values of 4 measurements + SD. Panel D shows typical plot of the time course of leaf weight changes in the dark. Data of panel e were calculated from the linear part of the curve indicated by the arrows. * and **, different from WT at P<0.05 and 0.01 respectively (Student’s t test)

The effect of *SMR4* and *SMR5* expression on drought tolerance was not saturated because application of β-CCA on the *SMR*-overexpressing lines increased further the resistance to water stress (Extended Data Fig. 5). The β-CCA signaling pathway is possibly wider than the effect of *SMR* overexpression. Indeed, β-CCA affects a larger panel of genes than the signaling pathway triggered in the OE:SMR5 plants (see below transcriptomic data). This can also be deduced from the effects of β-CCA on root length which was noticeably more marked than the effect of *SMR5* overexpression.

**Fig. 5.**
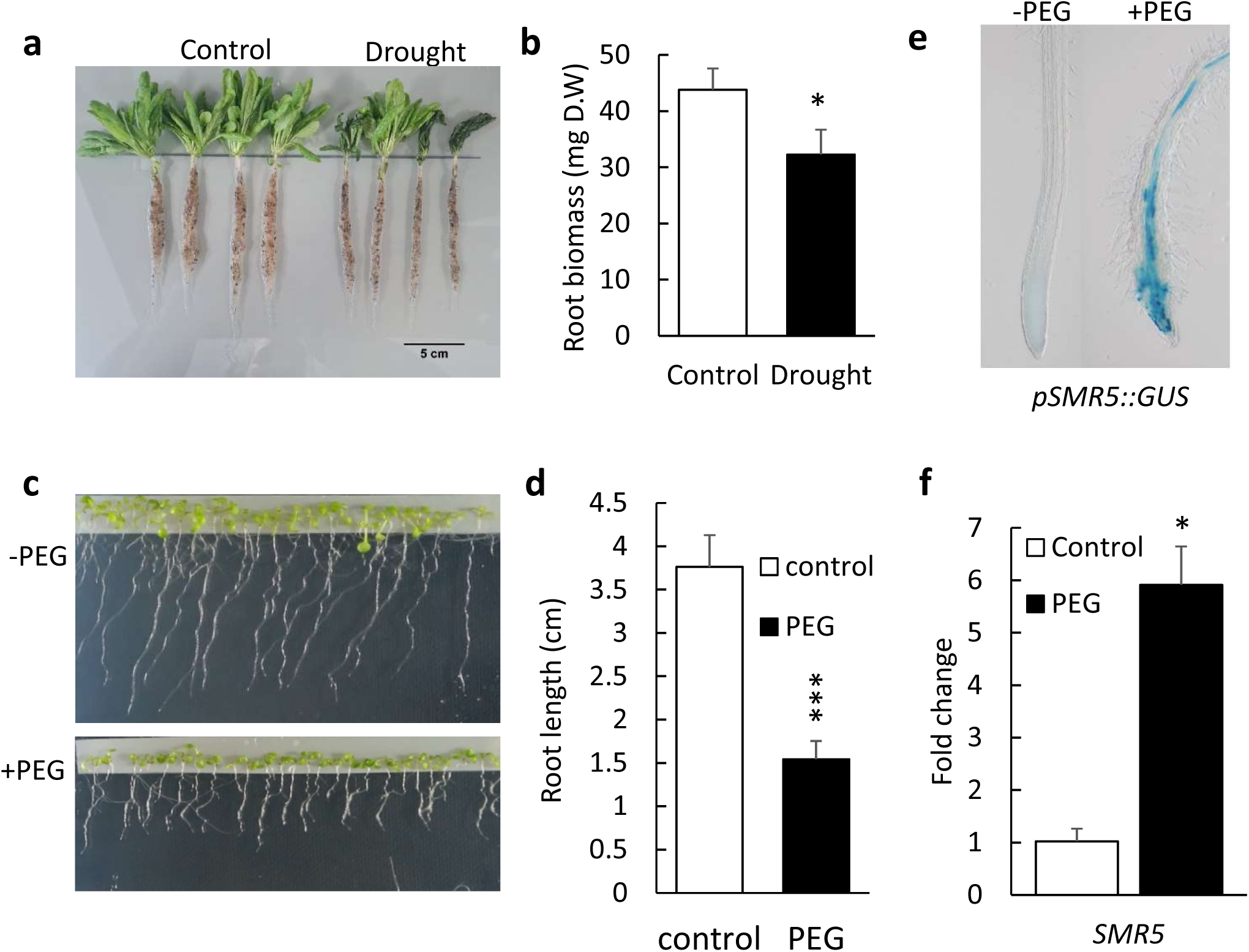
Effects of drought and osmotic stress on the Arabidopsis root system. A) Root system of control WT plants or drought-stressed plants grown on soil. B) Root dry weight. Values are means of 4 measurements + SD. *, P< 0.01 in two-tailed Student’s test. C-D) Root length of plants grown *in vitro* on Agar in the presence or absence of PEG 8000. Sedlings were grown on control medium for 3 d on Nylon stripes before transfer to PEG-enriched medium for 4 d. E) Expression pattern of SMR5 visualized by GUS coloration in the root tips of the GUS reporter line *pSMR5::GUS*. The plants were grown on Agar in the presence or absence of PEG 8000. F) qRT-PCR analysis of the transcript levels of *SMR5*. Data are mean values of 3 replicates +SD. They were normalized to the house keeping gene *UPL7* and to the control levels set to 1. * different from WT at P<0.05 (Student’s t test)

### SMR overexpression does not decrease stomatal conductance

Stomatal closure is a typical response of plants exposed to water stress conditions in order to limit water losses by transpiration (Munns 2002, Bartlett et al. 2016, Pirasteh-Anosheh et al. 2016). We analyzed the stomatal functioning in WT and *OE:SMR* plants using two complementary techniques, infrared (IR) imaging and porometry.

Leaf temperature varies with the transpiration rate: Opening of the stomata results in evaporative cooling and a decrease in leaf temperature, which can be monitored by thermal imaging (Jones 1999, Merlot et al. 2002). Fig. 4c shows images of plant temperature obtained with an IR camera. No difference was observed in leaf temperature between plants of the three genotypes under control conditions. Then, we exposed plants to a mild water stress induced by stopping watering for 4 d to avoid complete stomatal closure (like in Fig. 1a or Fig. 4a). Water stress induced a rise in leaf temperature reflecting a decrease in stomatal aperture in WT plants (Fig. 4c,d). The effect of drought was more marked in WT and *OE:SMR4* compared to *OE:SMR5*, likely due to the high drought tolerance of the latter genotype. Therefore the IR images support that stomatal transpiration is not reduced in the *OE:SMR* plants compared to WT plants, both under standard and stress conditions.

Porometry measurements were also conducted to evaluate the stomatal conductance by measuring the rate of water efflux from the leaf abaxial side. Under control conditions, stomatal conductance was similar in WT and *OE:SMR* plants (Fig. 4e), in agreement with Fig. 4c,d. Mild water stress conditions decreased stomatal conductance in WT and OE:SMR leaves, consistently with the increased temperature of leaves under those conditions. We can conclude that the increase in drought tolerance of *OE:SMR* plants does not rely on a reduction of stomatal aperture relative to WT.

Cuticular transpiration was measured by monitoring water losses from excised rosettes placed in the dark. Upon transfer of the plant rosettes from light to darkness, there was a rapid loss of water for ca. 30 min followed by a linear decrease in the plant mass (Fig. 4f). The first phase reflects the loss of water through the stomata which are closing in the dark (Hopper et al. 2014). The second phase reflects the cuticular losses of water after closure of the stomata. The second, linear phase was slowed down in *OE:SMR5* plants (-28%, Fig. 4f,g) and, to a lesser extent, in β-CCA-exposed plants compared to control WT rosette (-16%, Extended Data Fig. 6). This indicates a decrease in non-stomatal leaf transpiration by both β-CCA treatment and SMR5 overexpression.

**Fig 6.**
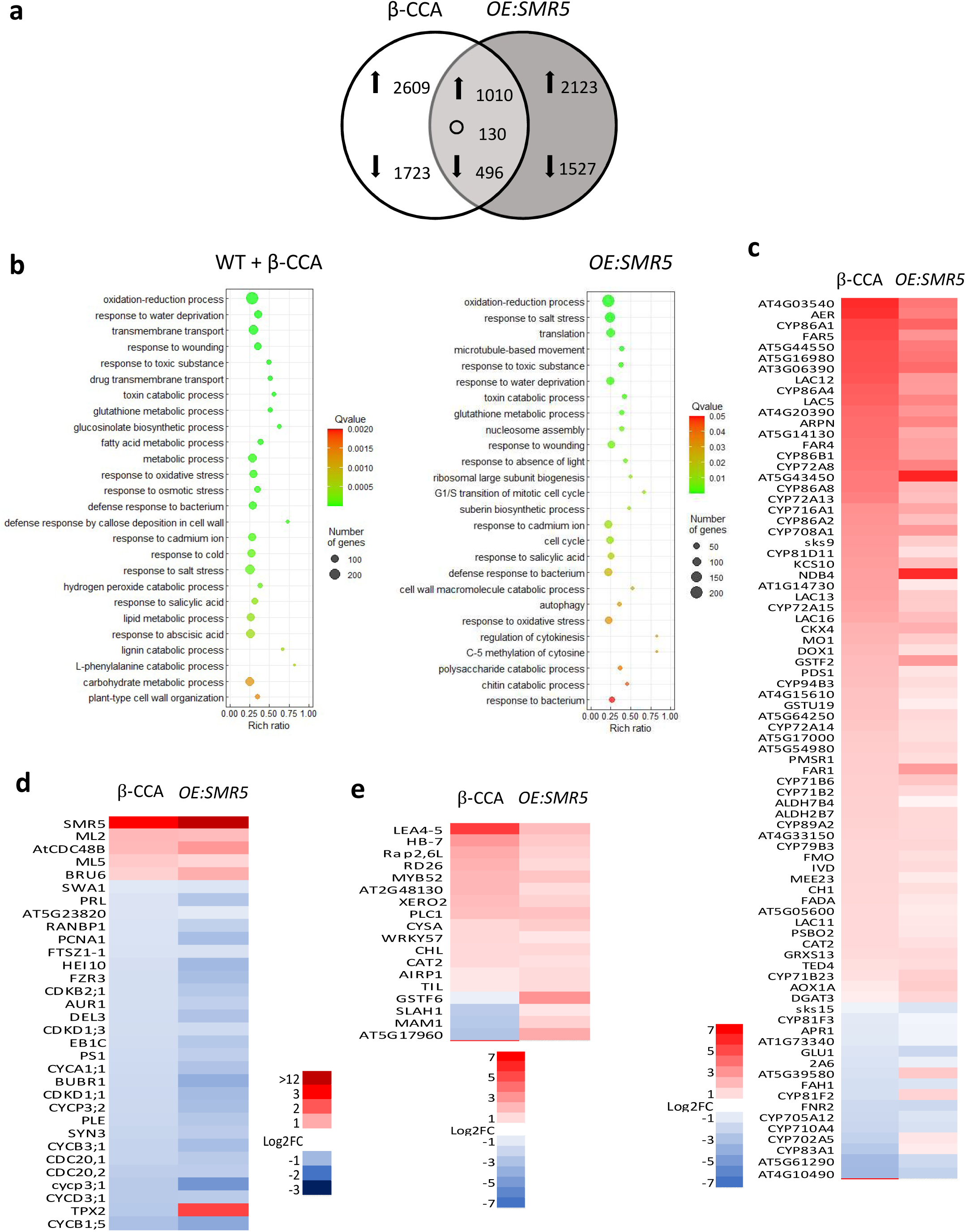
Transcriptome of Arabidopsis plants treated with β-CCA or overexpressing SMR5 compared to untreated WT plants. A) Venn diagram of up-regulated and down-regulated genes (P<0.05; log (fold change) ≥0) in roots of β-CCA treated seeding vs. untreated WT seedlings and of *OE:SMR5* seedlings vs. WT seedlings. Arrows indicate up-and down-regulation. O indicates counter-regulated genes. Data are mean values of 3 separate experiments. B) Gene ontology biological process (GOPB) enrichment bubble plots. Functional categories of the most affected genes by β-CCA or SMR5 overexpression are shown. Bubble color and size indicate the Q value and gene number enriched in the biological process. C-E) Heat maps of differentially expressed genes in roots of β-CCA-treated plants or plants overexpressing *SMR5* compared to untreated control WT plants. Genes are related to C) cell division, D) water deprivation, and E) oxido-reduction processes. P<0.05. Log (Fold Changes) ≥0.

### Water stress induces SMR expression and inhibits root growth

We previously showed that drought stress conditions induce β-CCA accumulation in Arabidopsis plants (D’Alessandro et al. 2019). Drought is also known to change root architecture (*e.g.* Verslues and Longkumer 2017, Kang et al. 2022), and this phenomenon is confirmed here by the reduction of root length in Arabidopsis plants exposed to drought compared to well watered conditions (Fig. 5a). Root dry weight was reduced by 25% in drought-stressed Arabidopsis plants (Fig. 5b). Similarly to plants grown on soil, Arabidopsis seedlings exposed to water deficit induced by adding PEG-8000 to the solid growth medium had shorter roots compared to control seedlings (Fig. 5c,d). This was associated with a substantial induction of *SMR5* (Fig. 5e,f). Thus, water stress is an environmental condition where the correlation between *SMR* induction and root growth inhibition found in β-CCA-treated plants is also observed.

### Spraying plants with a β-CCA solution does not have the same effect as watering plants with β-CCA

β-CCA can move within the plants through the xylemic flux: watering Arabidopsis plants with a solution of β-CCA was previously shown to lead to an accumulation of β-CCA in the leaves (D’Alessandro et al. 2019). Thus, we also measured the β-CCA concentration in roots of Arabidopsis plants grown on soil by GC-MS. In roots of control plants, the β-CCA concentration was low (<15 ng g^-1^ fresh weight), close to the detection limit of our GC system. This concentration is of the order of magnitude of the β-CC concentration in Arabidopsis roots (Dickinson et al. 2019). However, this is approximately 10 times lower than the β-CCA levels in leaves (139.53±16.05 ng g^-1^ fresh weight). In plants watered with 1.5 mM β-CCA, the β-CCA concentration in the roots drastically increased to 29.97±11.16 µg g^-1^ fresh weight. We cannot exclude that the low β-CCA concentrations measured in roots of soil-grown control Arabidopsis plants was due to β-CCA losses (or metabolization) during the rather long process of root washing. When the GC analyses were performed on young Arabidopsis seedlings grown *in vitro* on Agar, the measured concentrations in roots were noticeably higher, 864±263 ng g^-1^ fresh weight.

While β-CCA can move from the roots to the leaves (D’Alessandro et al. 2019), the opposite does not seem to occur: spraying Arabidopsis shoots with β-CCA (0.5 mM) did not result in an accumulation of the compound in the roots (14±5 and 12±2 ng g^-1^fresh weight in plants sprayed with water and β-CCA, respectively). Thus, β-CCA does not seem to be transported from the leaves to the roots by the phloem. Accordingly, spraying leaves with β-CCA did not inhibit root growth (Extended Data Fig. 7a,b) and did not enhance drought tolerance (Extended Data Fig. 7c). However, we cannot exclude that β-CCA is transported through the phloemic flux and subsequently metabolized in the roots.

**Fig. 7.**
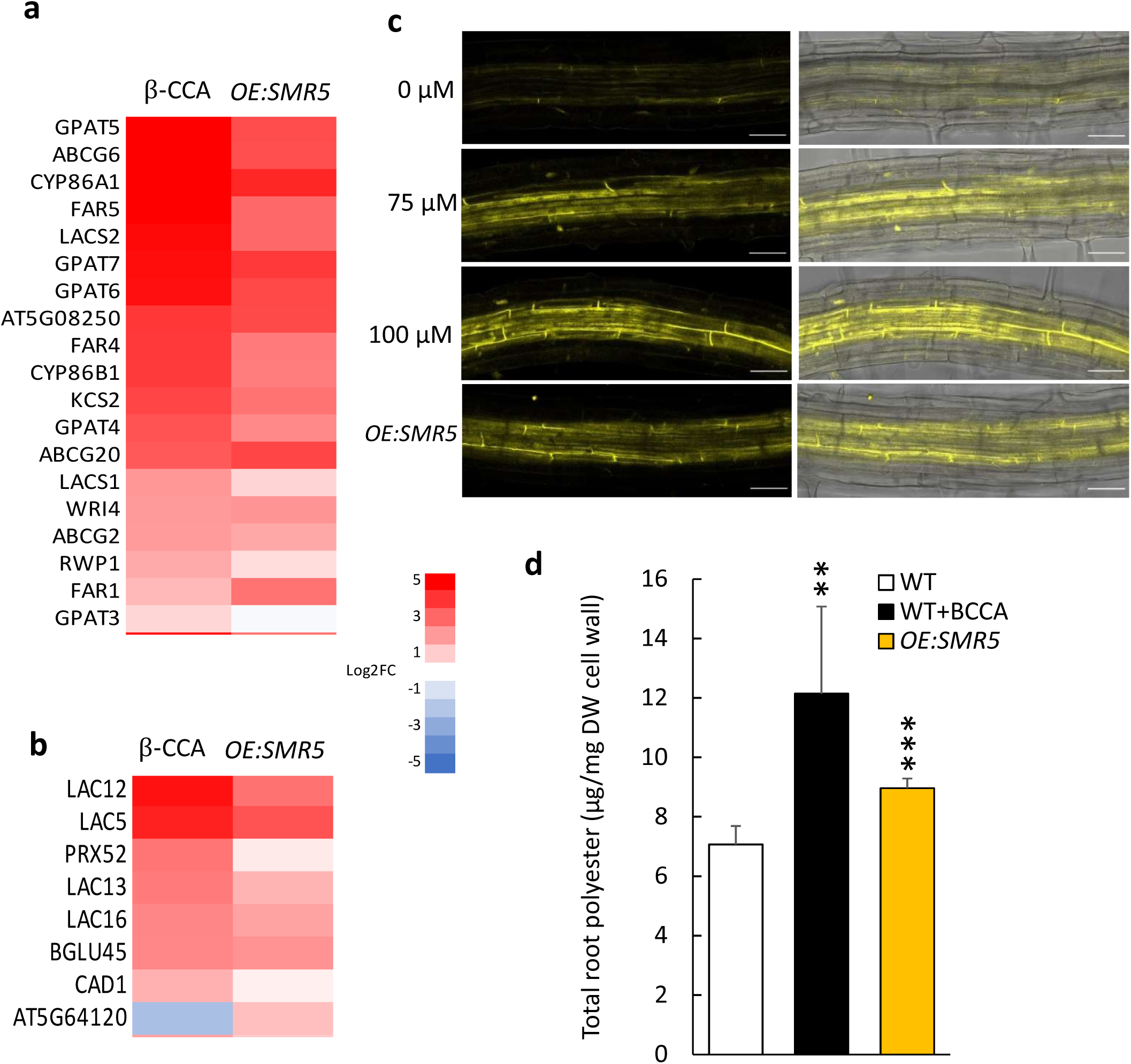
Root suberin in Arabidopsis plants treated with β-CCA or overexpressing SMR5. A) Heat maps of differentially expressed genes related to suberin and lignin biosynthesis. B) Suberin visualization by the fluorescence of Fluorol Yellow (left side). Scale bars, 50 µM. Right side: merged Fluorol Yellow fluorescence images and transmission light images. C) Total root polyesters. Data are mean values of 4 to 6 separate measurements + SD. *, different from WT at P < 0.05 (Student’s t test).

### Large overlaps between the transcriptomic responses to β-CCA treatment and to SMR5 overexpression

We performed a transcriptomic analysis of root tips from control and β-CCA-treated WT plants by RNA sequencing (RNA-seq). Around 4300 genes were differentially expressed between the two types of plants. The Venn diagram of Fig. 6a shows that β-CCA induced more than 2600 genes and decreased the expression of more than 1700 genes compared to untreated control plants. Classification of the differentially expressed genes in functional categories (Fig. 6b) revealed interesting features of the β-CCA effects on gene expression. The highest numbers of genes responsive to β-CCA were found in categories that are related to the responses to water deprivation/osmotic stress/oxidative stress, to oxidoreduction processes (including detoxification processes), to transmembrane transport, to cell wall and to different metabolic processes. Changes in gene expression in those functional classes fit with the idea that β-CCA induces a global response to water stress involving cellular detoxification and defense mechanisms and cell wall modifications.

RNA-seq analysis was also conducted on root tips of the SMR5 overexpressor. The number of genes induced or repressed compared to WT was above 2100 and 1500 genes, respectively (Fig. 6a). This is slightly lower than the number of genes induced/repressed by β-CCA (3600 vs. 4300). There was a substantial overlap in the transcriptomic responses to β-CCA and to SMR5 overexpression with about 1000 upregulated and 500 downregulated genes in common. Very interestingly, there was a noticeable concordance in the functional categories of the most affected genes by SMR5 overexpression and by β-CCA. Similarly to the transcriptomic signature of the β-CCA treatment, there was an enrichment in genes related to oxidoreduction processes, responses to water deprivation/osmotic stress/oxidative stress and cell wall in *OE:SMR5* vs. WT.

The heat maps of genes linked to cell division (Fig. 6c), water deprivation (Fig. 6d) and oxido-reduction processes (Fig. 6e) confirm the strong homology between the gene signature of β-CCA-treated plants and SMR5-overexpressing plants. Except a very limited number of genes (∼8 genes) that behaved in a opposite manner in the two conditions, all the genes induced by β-CCA were also induced by *OE:SMR5* and similarly all the genes downregulated by β-CCA were down-regulated by SMR5 overexpression in each gene category. Interestingly, a number of genes belonging to the ‘cell division’ category, including a number of cyclins and CDKs, were down-regulated in β-CCA-treated plants and *OE:SMR5*, consistently with the inhibitory effect on root cell division (Fig. 6c). The repression levels appeared to be higher in *OE:SMR5* than in β-CCA-treated plants. SMR5 was one of the most induced genes by β-CCA in this category. There was a high concordance in the induction/repression of genes related to oxidoreduction processes between β-CCA-treated plants and *OE:SMR5* (Fig. 6e). Finally, both β-CCA and SMR5 upregulation induced many genes responsive to water deprivation (Fig. 6d).

### Effect of β-CCA and SMR5 on root suberin and leaf cuticular lipids

The transcriptome of β-CCA-treated plants and *SMR5*-overexpressing plants revealed modifications of the expression of genes involved in the biosynthesis of the cell wall-associated biopolymers suberin and lignin (Fig. 7a,b) and in the deposition of callose in cell walls (Fig. 6b). Changes in those compounds can have significant effects on root and leaf hydraulics (Schreiber 2010). Among the genes known to be involved in suberin/cutin biosynthesis (Li-Beisson et al. 2013), LACS, several GPAT genes (GPAT 4, 5, 6, 7), several FAR, ABCG and CYP genes were noticeably up-regulated (Fig. 7a).

Suberin was imaged in roots using the fluorescent probe Fluorol Yellow (Marhavy and Siddique 2021). The intensity of the fluorescence signal was much stronger in roots of β-CCA-treated seedlings and in *OE:SMR5* seedings compared to WT (Fig. 7c). This suberization of the root tissues was confirmed by GC-MS analyses: total lipid polyesters of roots increased by a factor of almost 2 from 7.3 to 12.2 µg mg^-1^ cell wall after β-CCA treatment of the plants (Fig. 7d). Most polyesters were increased, but the strongest relative enhancement was observed for the 22:0 and 24:0 fatty acids, the 24:0 ω-OH and 18:1 ω-OH hydroxy fatty acids, and the 24:0, 18:1 and 18:2 α-ω dicarboxylic acids (DCAs) (Extended Data Fig. 8). The total lipid polyester levels were also increased in *OE:SMR5* roots relative to control WT (about +17%). The most enhanced monomers were the 18:1 and 16:0 ω-hydroxy fatty acids and 18:2 DCA (Extended Data Fig. 8).

It is possible that cuticular lipid polyesters are also upregulated in leaves by β-CCA. The decrease in leaf cuticular transpiration shown in Fig. 4 could suggest an increase of cuticular lipids. However, this was not observed when cutin and waxes were analyzed by GC-MS: No effect on the cutin levels was found in response to the β-CCA treatment or the *SMR5* overexpression (Extended Data Fig. 9). Wax coverages did not increase either in those plants (Extended Data Fig. 10).

## Discussion

### β-CCA is a root growth inhibitor

This work has shown that the apocarotenoid β-CCA, which accumulates in plants under abiotic stress conditions (D’Alessandro et al. 2019, Havaux 2020), has negative effects on root growth. This compound impacts root development by inhibiting both cell division and expansion. It is well established that carotenoid-derived phytohormones, such as abscisic acid and strigolactones, shape plant root architecture under specific environmental conditions (Creelman et al. 1990, Sharp et al. 1994, Koltai et al. 2011). Recently, a number of bioactive metabolites derived from ROS-mediated or enzymatically catalyzed oxidation of carotenoids, such as β-CC, retinal, (iso-)anchorene and zaxinone have been identified as new root growth regulators (Ke et al. 2022). β-CC, retinal, anchorene and zaxinone have been reported to act as stimulators of root growth (Dickinson et al. 2019 and 2021, Jia et al. 2019, Wang 2019) whereas iso-anchorene conversely inhibits primary root growth (Jia et al. 2021). β-CCA is another apocarotenoid that down-regulates root development, affecting both primary and secondary roots. This work thus reinforces the recent view that a network of carotenoid-derived metabolites plays essential roles in root development and in shaping root system architecture (Ke et al. 2022). Similarly to anchorene and its isomer iso-anchorene, β-CC and its oxidation product β-CCA appear to have opposite effects on root growth, suggesting that the in vivo role of the root apocarotenoid network could be rather complex. From a more general point of view, our results fit with the idea that modulating the content of carotenoids and of some of their derivatives could provide new tools to enhance plant yield and/or resilience (Moreno and Al-Babili 2022). Moreover, cell division and expansion at the root tip are known to be reduced when water availability drops (Sanchez-Calderon et al. 2013, Verslues and Longkumer 2017, Kang et al. 2022), as confirmed here by the inhibition of Arabidopsis root growth by water stress/osmotic stress conditions. Consequently, we can propose that the stress-induced, root-inhibiting β-CCA signal plays a role in the root growth response to drought.

### β-CCA inhibits the root cell division mechanism

This work provides clues on the mode of action of β-CCA on root growth and drought tolerance. The expression of components of the cell division mechanism, including several CDKs and CYCs, was lowered in root tips by β-CCA, whereas CDK-CYC inhibitors, especially *SMR5*, were up-regulated. The concomitant and opposite changes in root cell division activators (CDK and CYC) and inhibitors (*SMR*) by β-CCA are expected to negatively affect root growth. Accordingly, overexpression of a *SMR* gene (*SMR1*) and accumulation of the corresponding protein were previously reported to cause root growth inhibition (Dubois et al 2018). Interestingly, *SMR5* was shown to be the most sensitive *SMR* gene to moderate drought, displaying an increased expression in Arabidopsis leaves at all developmental stages (Dubois et al. 2018). A number of other *SMR* genes are also responsive to drought, but this was strongly dependent on leaf age and development. In maize, expression analyses of *SMR* genes indicated that they are strongly associated with abiotic stresses including salt stress and osmotic stress (Zhang et al. 2020). On the other hand, cyclin expression is a limiting factor for root growth, and overexpression of *CYC1* was found to accelerate root growth (Doerner et al. 1996). Conversely, inhibition of *CYCP3;1* expression by the brassinosteroid signaling was reported to inhibit root growth (Chen et al. 2020), and a mutant deficient in *CYCD4;1* exhibited a decreased density of lateral roots (Nieuwland et al. 2009). Reduction of root growth rate by salt stress was associated with a decrease in the kinase activity of CDKs (West et al. 2004), and loss of CDKC;2 increased drought tolerance in Arabidopsis (Zhao et al. 2017). Transcriptome analyses revealed that multiple *CYC* and *CDK* transcripts are downregulated after osmotic stress (Skirycz et al. 2011). Those previous observations indicate that the β-CCA-induced perturbations of the cell division machinery can not only inhibit root growth, but it can also lead to an enhancement of drought tolerance. Our results and previous observations (Yi et al 2014, Reyt et al. 2015, Dubois et al. 2018) also indicate that, compared to other *SMRs*, *SMR5* expression is particularly responsive to environmental stresses.

### SMR cell cycle inhibitors promote drought tolerance

Induction of *SMRs* by β-CCA appeared to be instrumental in the enhancement of drought tolerance by the apocarotenoid. Indeed, overexpression of these genes in transgenic Arabidopsis was sufficient to enhance plant tolerance to drought stress. *SMR5* overexpression also affected the expression of a large number of genes. Moreover, there was a strong overlap in the transcriptomic responses of Arabidopsis plants treated with β-CCA or over-expressing *SMR5*. Thus, *SMR5* expression partially mimics the effects of β-CCA in both gene expression changes and physiological responses, suggesting that *SMR5* is an upward component in the β-CCA signaling pathway. Interestingly and similarly to β-CCA (D’Alessandro et al. 2019), *SMR5* upregulation induced the expression of a number of drought stress marker genes (RAP2.6, RD26, LEA4-5, HB7, …). Thus, high *SMR* expression is somehow perceived by the plant as a drought stress signal. As mentioned above, *SMRs* are inhibitors of the CDK-CYC complexes and, in agreement with our results, downregulation of CYCs and CDKs have been previously associated with drought tolerance (Zhou et al. 2013, Zhao et al. 2017).

The transcriptomes of β-CCA-exposed roots and SMR5-overexpressing roots provide some indications on how plants became resistant to severe drought stress. In leaves, a major transcriptomic effect of β-CC and β-CCA was the induction of the xenobiotic detoxification pathway (D’Alessandro et al. 2018, 2019). This pathway detoxifies toxic and reactive compounds generated during stress, such as reactive carbonyl species generated from lipid peroxidation, by a network of detoxifying enzymes, limiting cellular injuries and restricting propagation of oxidative stress (Rac et al. 2021). Similar to β-CC(A)-treated leaves, many genes of the xenobiotic response (e.g. AER, two ALDH genes, several GSTU genes and many CYP genes encoding Cytochromes P450, …) were also induced in roots by β-CCA and SMR5. This effect can contribute to drought tolerance by preserving root cell integrity. Accordingly, preservation of the structural integrity of lipid membranes to avoid that they become leaky is believed to be one of the first line of defense against cellular dehydration in plants exposed to drought stress (Chng et al. 2022).

Spraying leaves with β-CCA did not bring about an accumulation of β-CCA in the roots. Also, root growth was not inhibited and drought tolerance was not enhanced after a spray of β-CCA. Therefore, root accumulation of β-CCA and modification of the root physiology seems to play a crucial role in the enhancement of drought tolerance by β-CCA. Higher drought tolerance of plants with shorter or less dense root system is somewhat counterintuitive; one can indeed imagine that root proliferation could increase the capture of water through better soil exploration. However, deep roots for water acquisition confer advantages for plants growing under conditions where deep stored water is available (Comas et al. 2013), obviously not for plants that must capture shallow water or that are grown in containers. In our experiments, we measured soil moisture at the top and the bottom of the pots at different stages of water stress (0, 7 and 12 d). The changes in soil moisture were very similar for water-stressed WT and *OE:SMR5* plants (Extended Data Fig. 11). Therefore, the high tolerance of *OE:SMR5* plants relative to WT cannot be attributed to a difference in soil drying during the stress experiment.

Terrestrial plants evolved in environments that favor phenotypes with aggressive root proliferation, but is has been argued that parsimonious root phenotypes with reduced crown root production and less lateral roots is advantageous for drought resistance in high-input agroecosystems (Lynch 2018). Van Oosterom et al. (2016) showed that maize genotypes capable of maintaining transpiration at low root mass show a better adaptation to drought stress. In Arabidopsis too, there are examples where drought tolerance is associated with reduced root development. Overexpression of the transcription factor MYB96 in Arabidopsis concomitantly inhibited lateral root elongation (Janiak et al. 2016) and promoted drought tolerance (Seo et al. 2008). The increased tolerance to drought of ATAF1-or RGS1-overexpressing Arabidopsis lines was also accompanied by an inhibition of root growth (Chen et al. 2006, Wu et al. 2009).

### β-CCA and SMR5 enhance waterproof barriers in roots

We previously showed that neither β-CC nor β-CCA modify stomatal behavior: treatments of plants with those compounds did not trigger stomatal closure (Ramel et al. 2012, D’Alessandro et al. 2019), and the response of stomata to water stress was unchanged compared to control, untreated plants (D’Alessandro et al. 2019). This is confirmed here by porometric measurements of stomatal conductance and by IR imaging of leaf temperature in β-CCA-treated plants. Similarly, SMR5 upregulation did not promote stomatal closure compared to WT, both in control conditions and during water stress. We can conclude that the increased tolerance to drought stress by β-CCA or SMR5 expression does not rely on changes in stomatal transpiration. However, plant leaves can also loose water via the cuticular transpiration. Although the rate of the latter mechanism is small compared to stomatal transpiration, the residual transpiration (by closed stomata and cuticle) can play a significant role under drought stress conditions. We observed here that both SMR5 and β-CCA lowered the non-stomatal water losses of leaves.

The transcriptomic changes induced by β-CCA and SMR5 revealed the induction of genes involved in several metabolic pathways leading to cell wall-associated biopolymers such as cutin, suberin or lignin. The leaf cuticle is the outermost layer of the leaf epidermis and is composed of a lipid polyester matrix (cutin) impregnated with waxes, a complex mixture of very-long-chain fatty acid derivatives and other hydrophobic compounds (Pollard et al. 2008, Yeats and Rose 2013). This lipidic ensemble constitutes a hydrophobic thin layer that can modulate nonstomatal water losses and thus contribute to drought tolerance (Xue et al. 2017, Monneveux et al. 2004, Petcu et al. 2009, Zhang et al. 2013, Xue et al. 2017). However, wax and cutin amounts are not always correlated with cuticular transpiration. In barley, osmotic stress increased leaf wax and cutin amounts without any change in cuticular conductance (Shellakkutti et al. 2022). In our study too, we could not correlate cutin and wax levels with the changes in cuticular transpiration. Therefore, other parameters of leaf hydraulics must play a role in lowering of nonstomatal transpiration by β-CCA and SMR5 expression. They may have led to the accumulation of other components of the cuticle, like e.g. the non-saponifiable cutan polymer (Boom et al. 2005) and/or to the establishment of internal barriers regulating the flow of water from the leaf tissues to the atmosphere (Geng et al. 2022, Prado and Maurel 2013). Interestingly, a number of aquaporin genes were found to be downregulated in the leaf transcriptome of Arabidopsis plants exposed to volatile β-CC: PIP1;1 (AT3G61430), PIP1;2 (AT2G45960), PIP2;2 (AT2G37170) and PIP1;5 (AT4G23400) (Ramel et al. 2012). Because a substantial fraction of β-CC is converted to β-CCA in planta (D’Alessandro et al. 2019), the β-CC-induced transcriptome may be, at least partially, a β-CCA transcriptome. Therefore, decreased expression of aquaporin genes may be a factor that could change water transport in β-CCA-treated leaves. The exact causes of the decrease in non-stomatal transpiration in the absence of a significant change in the leaf cuticle remain, however, to be determined.

β-CCA-exposed roots exhibited high levels of suberin monomers compared to control WT plants. Suberin is a lipophilic polyester associated to the cell walls in some tissues such as the endodermis and the periderm of roots, and functions as a diffusion barrier for water, gases and solutes (Franke et al. 2012, Shukla and Barberon 2021, Woolfson et al. 2022, Serra and Geldner 2022). Suberin deposition in roots is enhanced under environmental stress conditions including drought stress, decreasing root hydraulic conductivity (Franke at al. 2012, Karlova et al. 2021). Arabidopsis mutants with increased root suberization exhibited lower root hydraulic conductivity, lower transpiration rate and increased tolerance to drought compared to WT (Baxter et al. 2009). Consequently, suberin accumulation in Arabidopsis roots by β-CCA and SMR5 upregulation could participate in the enhancement of drought tolerance by limiting water losses during water stress. Interestingly, the transcriptomic data reveal β-CCA-and SMR5-induced changes in the expression of genes related to the biosynthesis of other biopolymers than cutin/suberin, such as callose and lignin, which could further reinforce this protective mechanism against drought stress (Moura et al. 2010, Cui and Lee 2016, Wang et al. 2022). Increased lignin levels in roots have been previously associated with enhanced tolerance to drought (Li et al. 2021). Callose deposition in plants is mainly considered as a component of the immune system, limiting pathogen invasion and spread (Wang et al. 2022). However, there are some reports correlating increases in callose levels with the response and/or tolerance to abiotic stresses (e.g. Pilai et al. 2018). It thus appears that β-CCA promotes a general upregulation of defense mechanisms that provide water-holding protection of roots and leaves, maintaining water content of plant tissues during stress exposure.

## Conclusions

The reprogramming of gene expression by β-CCA impacts a large panel of genes. One can then expect that the biological effect of β-CCA on plant drought tolerance is a multifactorial process involving various aspects of the plant functioning. Here, we have identified a novel element in β-CCA signaling with major roles in drought tolerance. β-CCA triggered the induction of cell cycle inhibitors of the SMR family, and the selective upregulation of SMR5 in absence of exogenous β-CCA led to a remarkable enhancement of drought tolerance and to changes in gene expression that resemble very much those induced by β-CCA. The highly overlapping responses between SMR5 overexpression and the β-CCA treatment would position SMR5 upstream of the β-CCA/drought tolerance pathway. Based on our results, SMR5 constitutes a new target for improving plant tolerance to water stress. Although SMRs are inhibitors of cell division, overexpression of SMR5 or SMR4 had modest effects on shoot growth. In contrast, the effect on roots was much more pronounced, although dependent on the substrate on which the plants have grown. Similarly, growth of the aerial parts of Arabidopsis plants was not affected by β-CCA (D’Alessandro et al. 2019) whereas roots were noticeably impacted (this study). Because of the marked effects on roots, modulation of SMR5 levels to enhance drought tolerance would probably be most useful in crop plants that are exploited for their leaves and/or fruits, rather than for underground parts.

It appears from our results that reorientation of root metabolism from growth and development towards defense mechanisms is an important component of drought tolerance of Arabidopsis and of the physiological response to β-CCA. The present study has identified SMR5 and its biochemical inducer, β-CCA, as regulators of this phenomenon, hence providing new actors in the process by which plants can balance growth and stress response. Future strategies for resetting the balance between stress resistance and growth to engineer stress-resistant and high-yielding crops require the understanding of how stress signaling regulates plant growth (Zhang et al. 2020). Apocarotenoid-triggered induction of SMRs appears to be a piece of the puzzle which links the plant cell division machinery with the defense mechanisms against stress. Finally, a major challenge for future research in this area will be to identify the primary target/receptor of β-CCA that triggers the up-regulation of SMR5 expression.

## Methods

### Plant material and growth conditions

Wild-type (WT) *Arabidopsis thaliana* (ecotype Col-0), a triple SMR knockout mutant (*smr4 smr5 smr7*) and transgenic lines overexpressing *SMR5* or *SMR4* (*OE:SMR5* and *OE:SMR4*, respectively), provided by L. De Veylder (VIB, Belgium), were used in this study. A number of additional *OE:SMR5* and *OE:SMR4* transgenic lines have also been produced in the frame of this study (see below). Plants were grown on potting soil (Seedlingsubstrat, Klasmann-Deilmann) for 5 or 6 weeks in short-day conditions (8h/16h, light/dark) under controlled environmental conditions in phytotrons of the Phytotec platform (BIAM, CEA/Cadarache): the photon flux density was around 150 µmol photons m^-2^s^-1^, the air temperature was 20°C/18°C (day/night) and the relative air humidity was maintained at ca. 65 %. The size of the pots (one plant per pot) was 5.5 cm x 5.5 cm x 5 cm.

For *in vitro* cultures on 0.8% (w/v) Agar (Sigma Aldrich) in Petri dishes, Arabidopsis seeds were surface-sterilized for 2 min in a solution containing 70% (v/v) ethanol and 0.05% (v/v) sodium dodecyl sulfate, and washed twice with 95% ethanol. Seeds were then grown on vertical plates on 10 times diluted Murashige and Skoog medium (MS/10) under long-day conditions (16 h/8 h, day/night) at 22°C. The medium was supplemented with 2.5 mL of 0.20 µm-filtered 1.5 mM solution of β-CCA (2.5 ml in 50 ml of liquid medium, corresponding to a final concentration of 75 µM β-CCA, unless specified otherwise) or with ultrapure water (for the controls). The β-CCA solution was produced as previously described (D’Alessandro et al. 2019). In brief, 1 mL of pure β-cyclocitral (Sigma-Aldrich) was injected in 1 L of pure water and stirred vigorously for 24 h in a closed container. Then, the container was opened to eliminate volatile β-cyclocitral molecules that were not oxidized while continuing to stir vigorously for 24 h. The final β-CCA concentration, measured by GC-MS (see below), was around 1.5 mM β-CCA. Primary root length of Arabidopsis seedlings grown *in vitro* was measured after 6 d with the ImageJ software (NIH) using the NeuronJ plugin. Roots of plants grown on soil or on sand were carefully washed in water before measuring root length and dry weight. In most experiments, β-CCA was applied to the plants through the roots by watering the pots with a 1.5 mM solution of β-CCA (25 ml per plant in pots of about 150 cm^3^). In some experiments, plants were also sprayed with 0.5 mM β-CCA or with water for the controls.

For hypocotyl length measurements, the sterilized seeds were sown on Petri dishes containing the MS/10 medium supplemented with different concentrations of β-CCA. They were put at 4°C overnight, and then transferred to the dark at 24°C for 7 d. The length of the hypocotyls was measured with ImageJ.

### β-cyclocitric acid analyses

1 mL of 5 % H_2_SO_4_ (v/v) in methanol is added to 50 mg of plant tissue and heated at 90°C for 90 min. The sample is cooled down to room temperature and vortexed. 500 µL of hexane is then added, followed by 1.5 mL 0.9% NaCl (w/v). The mixture is vigorosly stirred for 40 s with a vortex mixer and centrifuge for 2 min. The upper phase is analyzed by GC-MS (7890A gas chromatograph and 5975C mass spectrometer; Agilent Technologies) using a polar OPTIMA WAX column (30 m 3 0.25 mm 3 0.5 mm, Macherey-Nagel). The GC conditions are as followed : splitless mode injection; injector temperature 240°C; oven temperature program: 50°C for 1 min followed by a temperature ramp of 10°C min^-1^ to 300°C, holding this temperature for 2 min. The flow rate of the carrier gas (He) was 1 mL min^-1^. Mass spectrometer paramater are positive mode and single ion monitoring mode. β-CCA was quantified on ions m/z 123 and m/z 135.

### Water stress treatments

Water withdrawal was applied to 5 week-old plants by stopping watering for 10 to 14 d. The relative water content of the leaves (RWC) was measured by weighing leaf disks (fresh weight, FW). The leaf disks were then fully hydrated for 24 h in water and in the dark (turgid weight, TW). Finally, the disks were dried at 70°C for at least 48 h (dry weight, DW). The RWC (in %) was calculated as [(FW-DW)/(TW-DW)] x 100. Soil moisture was measured by weighing a soil aliquot before and after drying in an oven for 48 h (F.W. and D.W., respectively). Soil moisture (in %) was calculated as [(F.W.-D.W.)/F.W.] x100.

Osmotic stress was applied to seedlings grown *in vitro* as described in van der Weele et al. (2000) and Verslues and Bray (2004). Polyethylene glycol (PEG 8000)-infused medium was prepared by overlaying 60 mL of PEG solution (200 g dissolved in 1 L MS/10) on 40 mL of solidified growth medium for at least 12 h. Control plates were overlayed with MS/10. The excess liquid solution was then poured off, and 3 d-old Arabidopsis seedlings were transferred on the Agar medium using sterile nylon strips.

### Stomatal conductance

Stomatal conductance of fully expanded leaves was measured on 6-week-old plants using a hand-held AP4 porometer (Delta-T Devices). Measurements were carried out on the abaxial leaf side in the middle of the light period following the instruction manual of the porometer. The apparatus was let to equilibrate in the phytotron for 2 h before measurements. A minimum of 10 measurements per plant were made on 6 plants per genotype.

Plant transpiration was also estimated by infrared (IR) imaging. Low relative humidity (45% ± 5%) and low wind speed were applied the day before IR thermographic imaging to ensure optimal contrast between lines. Images were acquired using a FLIR infrared camera of the A600 series, as described by Costa et al. (2015). The pixel resolution of the detector was 640 x 480, and the spectral range was 7.5-14 µm. For each comparison between WT and mutant lines, three independent experiments with at least four plants per genotype were carried out and led to similar results.

### Cuticular transpiration

Freshly cut Arabidopsis rosettes were immediately sealed with vacuum grease on the cut root collar. The rosettes were then placed on a tripod on the weighing pan of a precision balance and was let to dehydrate in complete darkness. The rosette weight was automatically measured every 3 min. At the end of the experiment, the plant was placed in an oven at 70°C to determine the dry weight.

### Microscopic analyses

Roots were stained with 0.01% (w/v) polycationic stain Ruthenium Red (Sigma Aldrich) for 10 min and observed with an Axio Zoom V-16 (Zeiss) microscope.

Confocal laser scanning microscopy imaging was performed with a Zeiss LSM780 or LSM980 confocal microscope. Excitation and detection windows were set at 514 nm and 650-700 nm, respectively, for Propidium Iodide, and 488 nm and 500-550 nm for Fluorol Yellow. Fluorol Yellow staining was performed as described in Leal et al. (2022). Roots were incubated in a freshly prepared solution of 0.01 % (w/v) Fluorol Yellow (Santa Cruz) in lactic acid (85%) at 70°C for 30 min. Afterwards, plants were washed three times in water. For Propidium Iodide staining, roots were incubated in a fresh solution of 10 µg L^-1^ for 5 min and then rinsed in water for 10 min.

Arabidopsis lines expressing cell cycle-related genes with the GUS marker (Barrada et al. 2019) were received from M.-H. Montane and B. Menand (BIAM, Marseille). The following homozygous GUS lines were used: a) a cyclin (CYC) gene fused with the GUS reporter for the translational reporter lines *CYCB1;1-GUS, CYCB1;2-GUS, CYCB3;1-GUS, CYCA2;3-GUS, CYCA3;1-GUS, CYCA3;2-GUS;* b) the promoter of *CYC-D/CCS52A1/SIAMESE/SIAMESE-RELATED* genes fused to the GUS marker for the transcriptional reporter lines *pCYCD3;3::GUS, pCYCD6;1::GUS, pCCS52A1::GUS, pSIM::GUS, pSMR4::GUS, pSMR5::GUS and pSMR7::GUS*. Seeds were grown in Petri dishes containing MS/10 medium. After 6 d, the seedlings were incubated at 37°C with X-glucuronidase solution (1mg/mL) containing 10 mM phosphate (pH 7), 10% (w/v) Triton X-100, 1 M potassium ferricyanide and 100 mM potassium ferrocyanide for 4h for *pSMR4::GUS, pSMR5::GUS and pSMR7::GUS* or overnight for the other lines. The staining was stopped with 75% (v/v) ethanol and mounted between slides and coverslips. GUS expression was observed with an Axio Zoom V16 (Zeiss) microscope.

### qRT-PCR

Total RNA was extracted using Direct-Zol RNA MiniPrep (Zymo Research) and treated with the RNase-free DNase Set (Zymo Research) according to the manufacturer’s instructions. Quantity of RNA was assessed using a NanoDrop2000 (Thermo Scientific, USA). Reverse transcription was performed on 400 ng of total RNA using qScript cDNA SuperMix (Quantabio). Quantitative PCR was performed on a 480 Light Cycler thermocycler (Roche) using the manufacturer’s instructions, using SYBR Green I Master (Roche) with 10 µM primers and 0.125 µL of RT reaction product in a total volume of 5 µL per reaction. The qPCR program was : 95 °C for 10 min, then 45 cycles of 95 °C for 10 s, 60 °C for 10 s and 72 °C for 10 s. The specific primers for PCR amplification, designed using the NCBI Primer designing tool, are listed in Extended Data Table 1. *UPL7* was used as reference gene for data normalization.

### Plasmid construction and plant transformation

Supplementary lines of *SMR* overexpressors were constructed. We received *E. coli* strains containing *SMR* open reading frames cloned into the pDONR221 entry vector by BP recombination cloning from Dr. Lieven De Veylder (VIB Belgium). The SMR open reading frames were then transferred into the pB2GW7 destination vector by recombinational cloning with Plant Gateway vectors (Karimi et al. 2007). All constructs were transferred into the *Agrobacterium tumefaciens* C58C1RifR strains. Transgenic plants in the T_2_ generation with T-DNA insertion at a single locus were selected by phosphinotricin resistance (10 mg L^-1^), and T_3_ homozygotes were used for all analyses. *SMR* gene expression was quantified in the overexpressing lines by qRT-PCR (see above) using *UCP* (Extended Data Table 1) as internal control for data normalization.

### RNAseq

Root tips (5-mm length) were harvested from Arabidopsis seedlings grown *in vitro* for 6 d. Total RNA was extracted by using Direct-zol RNA MiniPrep Plus (Zymo research, R2072). Quantity and quality of RNA were assessed respectively using NanoDrop2000 (Thermo Scientific) and Qubit RNA IQ assay kit (Thermo Scientific). Samples were analyzed by BGI Genomics (Hong Kong), providing a stranded mRNA library, 20M reads/sample, 100 bp paired end reads (100PE) on DNBseq^TM^. Standard bioinformatics analyses were performed by the Doctor Tom platform of BGI Genomics.

### Analysis of polyester monomers and cuticular waxes

Rosette leaves (250 mg fresh weight) of 4 week-old plants grown on soil were dipped in chloroform for 30 s and the cuticular waxes analyzed as previously described (Jakobson et al. 2016). Leaves were previously photographed to measure total area. For root polyester analysis, 100 mg (fresh weight) of roots were harvested from seedlings grown for 8 d on solid Agar medium and added to boiling isopropanol (85°C) containing 0.01% (w/v) butylated hydroxytoluene and heated for 10 min. For leaf polyester analysis, the same isopropanol procedure was applied to the leaves used for the wax analysis immediately after the chloroform remaining on the leaves had evaporated. Leaf and root cell wall polyesters were delipidated and depolymerized as previously described in detail (Li-Beisson et al. 2013). Briefly, the plant material quenched in isopropanol was delipidated and the cell wall polyesters were broken down to fatty acid methyl esters (FAMEs) using sulfuric acid-catalyzed depolymerization, the FAMEs were derivatized by acetylation and separated and quantified by gas chromatography coupled to mass spectrometry (GC-MS). GC-MS analysis was performed using conditions previously described (Jakobson et al. 2016).

## Data availability

The data generated in this study are provided in the Supplementary Information or from the corresponding author upon reasonable request.

## Acknowledgements

We would like to thank Lieven De Veylder (VIB, Belgium) for the kind gift of seeds of Arabidopsis SMR mutants and overexpressors and of *E. coli* entry clones with SMR ORF, Marie-Hélène Montane and Benoît Menand (BIAM) for providing seeds of Arabidopsis GUS lines, and Bernard Genty (BIAM) for the loan of a portable porometer. Useful discussions with Nathalie Leonhardt, Thierry Desnos and Laurent Nussaume (BIAM) are also acknowledged. Thanks are also due to Hélène Jacquet and Serge Chiarenza (BIAM) for technical advice, the Phytotec platform (BIAM) for their help in growing Arabidopsis plants and the Heliobiotec platform (BIAM) for making GC-and LC-MS equipment available to this study. This work was financially supported by ANR (ApoStress project) and PRIMA (UToPIQ project, N°1570).

## Author contributions

All authors participated in the scientific discussion. M.H. conceived and supervised the research. J.B. and M.H. designed the experiments. J.B. performed most experiments. M.L. and B. L. measured apocarotenoid contents. M.J. analyzed root growth characteristics. P.D. participated in the generation of transgenic plants. F.B. performed the analyses of lipid polysesters and waxes. S.D. and J.B. conducted bioinformatics analyses. J.B. and M.H. wrote the manuscript. All authors commented on the manuscript.

## Extended Data

Fig. 1. GUS coloration of root tips of the translational GUS reporter lines *CYCB1;1-GUS, CYCB1;2-GUS, CYCB3;1-GUS, CYCA2;3-GUS, CYCA3;1-GUS, CYCA3;2-GUS and the* transcriptional reporter lines *pCYCD3;3::GUS, pCYCD6;1::GUS, pCCS52A1::GUS, pSIM::GUS.*

Fig. 2. Root growth is not affected in the triple *SMR* mutant *smr4 smr5 smr7*.

Fig. 3. Growth curves of WT, *OE:SMR5* and *OE:SMR4* Arabidopsis plants.

Fig. 4. Drought tolerance of a series of *OE:SMR5* and *OE:SMR4* transgenic lines.

Fig. 5. Effect of β-CCA on the drought tolerance of *OE:SMR4* and *OE:SMR5*.

Fig. 6. Effect of β-CCA on leaf cuticular transpiration.

Fig. 7. Spraying plants with β-CCA does not induce the same response as watering plants with β-CCA.

Fig. 8. Profile of root suberin monomers in WT, 75 µM β-CCA-treated WT and *OE:SMR5* seedlings.

Fig. 9. Leaf cuticle in Arabidopsis plants treated with 1.5 mM β-CCA or overexpressing SMR5.

Fig. 10. Leaf waxes in Arabidopsis plants treated with 1.5 mM β-CCA or overexpressing SMR5.

Fig. 11. Soil moisture during the water stress treatment of WT and *OE:SMR5* plants.

