## Supplemental Figures and Table 1 for "An SMR cell-cycle inhibitor inducible by a carotenoid metabolite resets root development and drought tolerance in Arabidopsis"

### Extended Data

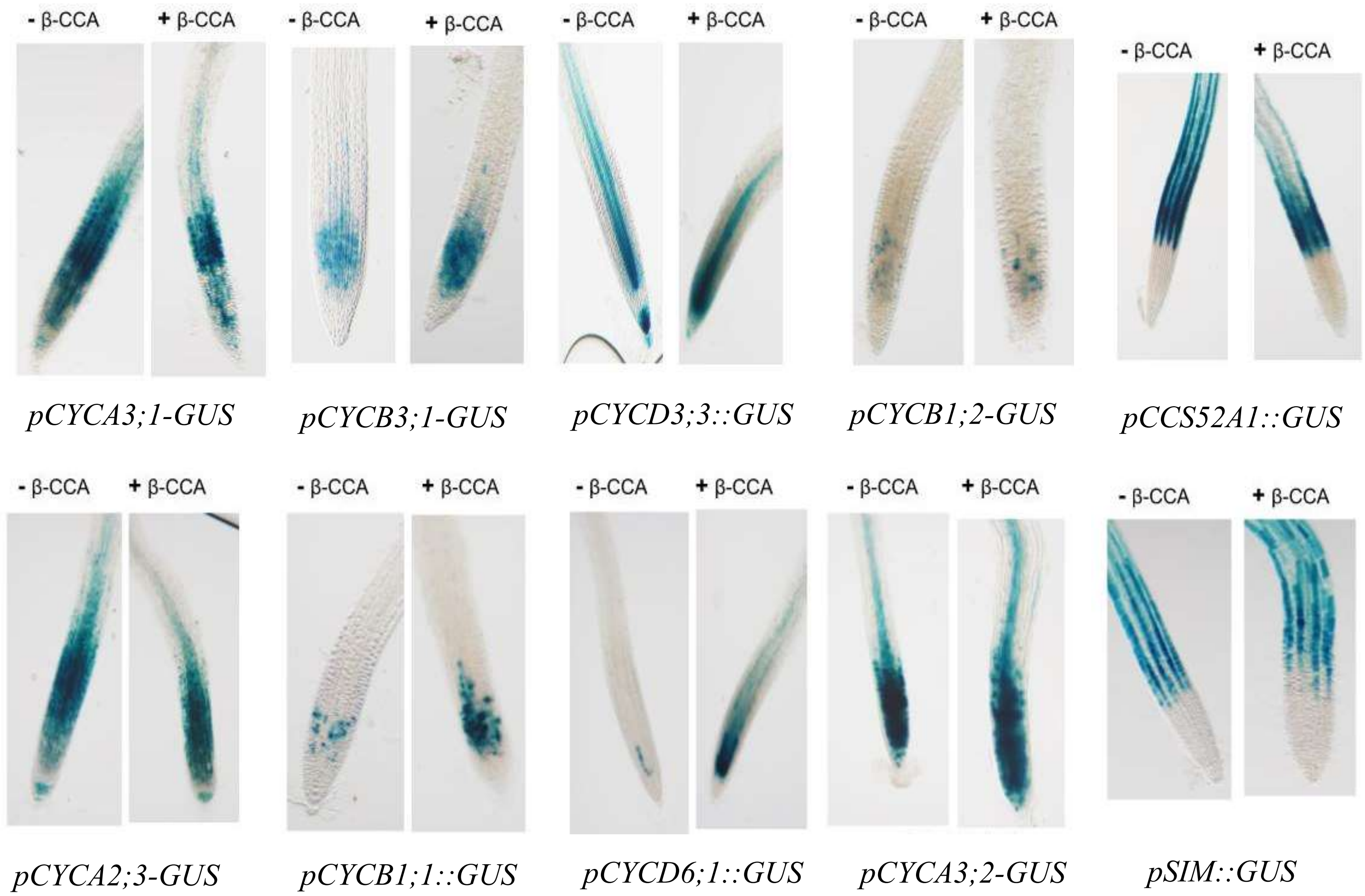


**Extended Data Fig. 1**. GUS coloration of root tips of the translational GUS reporter lines *CYCB1;1-GUS*, *CYCB1;2-GUS*, *CYCB3;1-GUS*, *CYCA2;3-GUS*, *CYCA3;1-GUS, CYCA3;2-GUS* and the transcriptional reporter lines *pCYCD3;3::GUS, pCYCD6;1::GUS*, *pCCS52A1::GUS*, *pSIM::GUS*. Seedlings were exposed to 0 or 75 µM β-CCA in the growth medium.

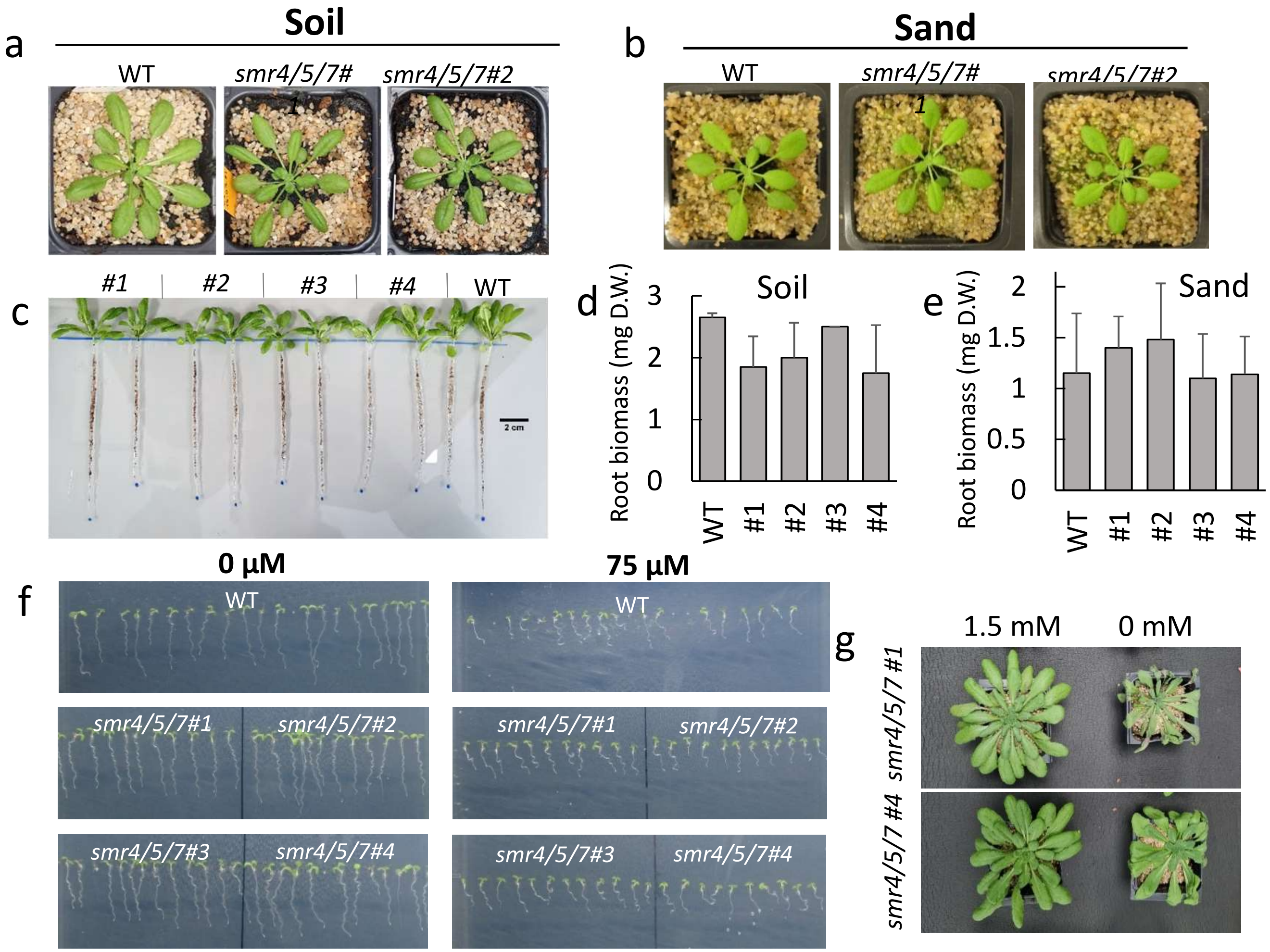


**Extended Data Fig. 2.** Root growth is not affected in the SMR triple mutant *smr4 smr5 smr7*. Representative rosettes of *smr4 smr5 smr7* triple mutants grown a) on soil for 5 weeks or b) on sand for 4 weeks. c) Root system of plants grown on soil for 5 weeks. d) and e) Roots were thoroughly washed under running water and then dried to measure dry weight (D.W.). Values are means + SD (n=2) in d) (n=5) in e). f), WT and mutants were grown *in vitro* with 0 or 75 µM β-CCA to check root length. g) Drought response of soil-grown triple *smr* mutants pretreated with 0 or 1.5 mM β-CCA.

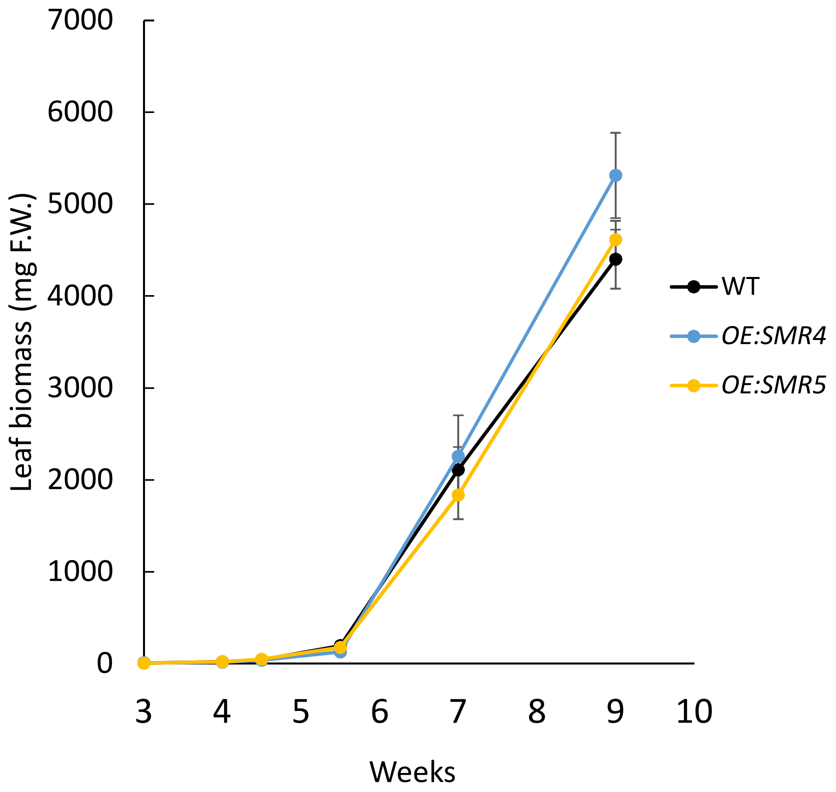


**Extended Data Fig. 3.** Growth curves of soil-grown WT, *OE:SMR5* and *OE:SMR4* Arabidopsis plants.

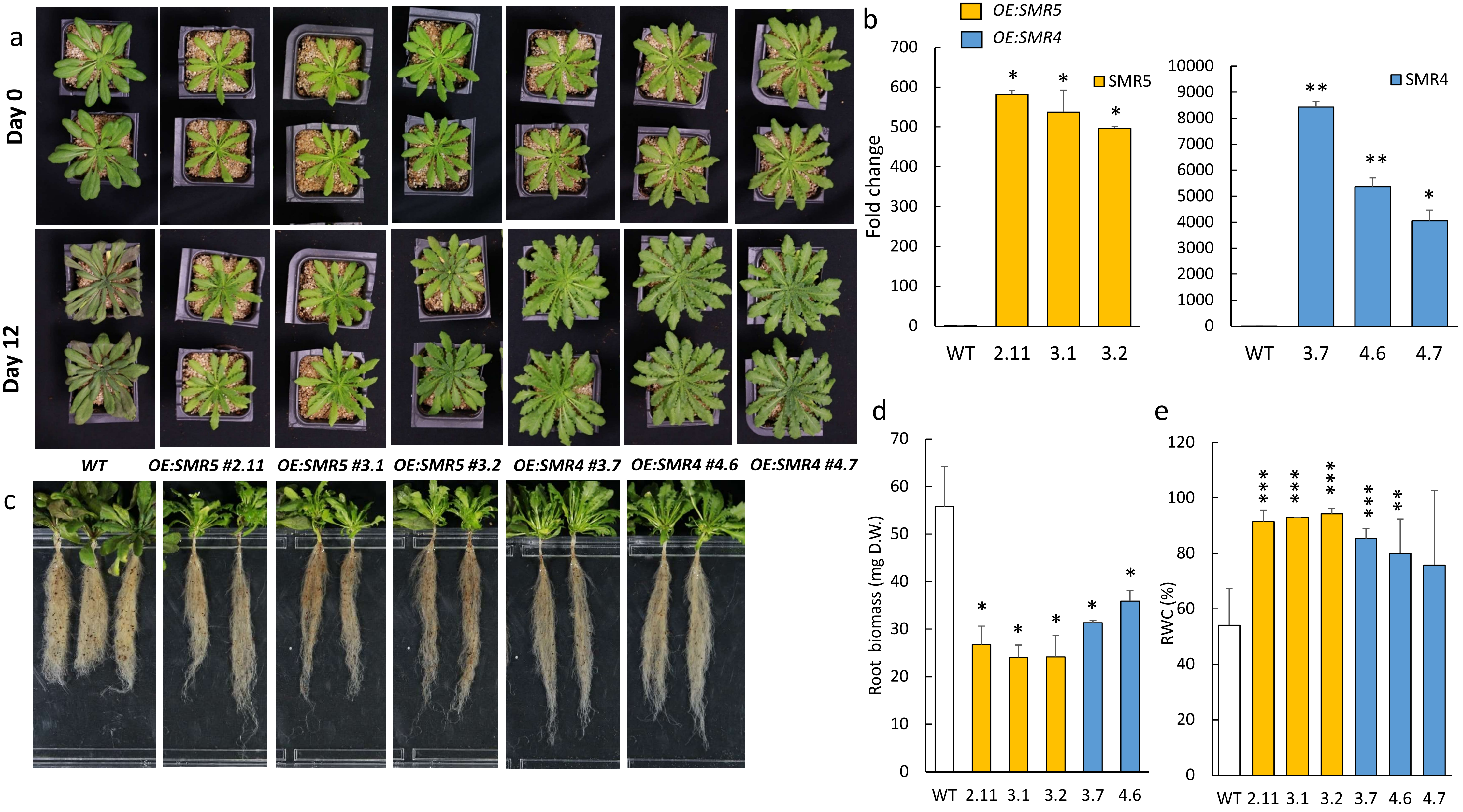
a
Day 0
Day 12
WT
OE:SMR5 #2.11
OE:SMR5 #3.1
OE:SMR5 #3.2
OE:SMR4 #3.7
OE:SMR4 #4.6
OE:SMR4 #4.7
b
OE:SMR5
OE:SMR4
Fold change
SMR5
SMR4
WT 2.11 3.1 3.2
WT 3.7 4.6 4.7
c
d
Root biomass (mg D.W.)
WT 2.11 3.1 3.2 3.7 4.6
e
RWC (%)
WT 2.11 3.1 3.2 3.7 4.6 4.7

**Extended Data Fig. 4.** Drought response of a series of independent *SMR4*- and *SMR5*-overexpressing transgenic lines**.** a) Picture of *OE:SMR4* and *OE:SMR5* lines before and after 12 d of water deprivation. b) Expression levels of *SMR4* and *SMR5* genes in the respective *OE:SMR* lines. c) and d) Pictures and dry weight of the root system of WT and *OE:SMR* lines after 12 d of drought stress. Data are mean values of 3 experiments + SD. e) Leaf RWC of plants after 12 d of water deprivation. Data are mean values of at least 3 experiments + SD. *, **, ***, different from WT at $P < 0.05$, 0.01 and 0.001, respectively (Student's t-test).

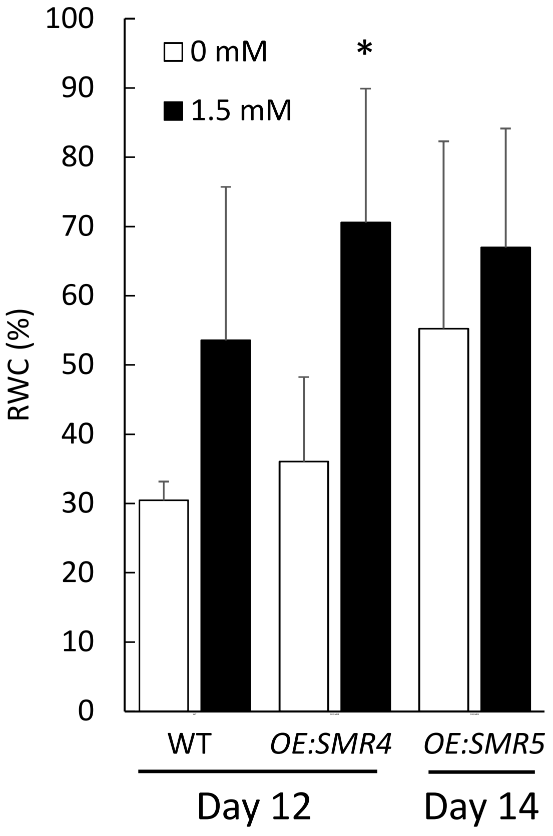


**Extended Data Fig. 5.** Effect of 1.5 mM β-CCA on the drought tolerance of WT, *OE:SMR4* and *OE:SMR5* plants, as measured by the leaf RWC. *, different from 0 mM at $P < 0.05$ (Student's t-test), $n$=2 to 4.

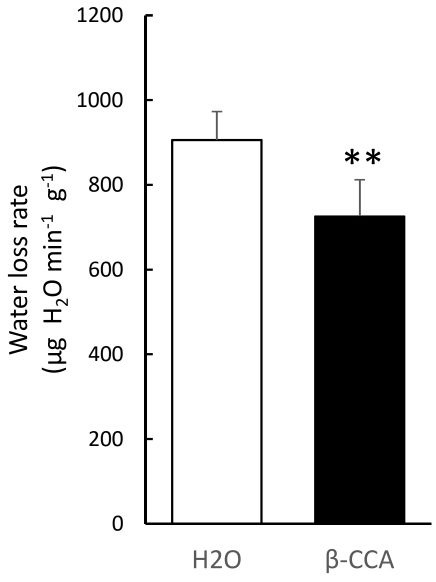


**Extended Data Fig. 6**. Effect of 1.5 mM β-CCA on leaf cuticular transpiration. Plants were pre-treated with 0 or 1.5 mM β-CCA for 4 d, then rosettes were excised, placed in the dark, weighed at regular intervals of time to measure water losses. Cuticular transpiration was calculated as in Fig. 4 f,g. **, different from $H_2O$ at $P < 0.01$ (Student's t test), n=4.

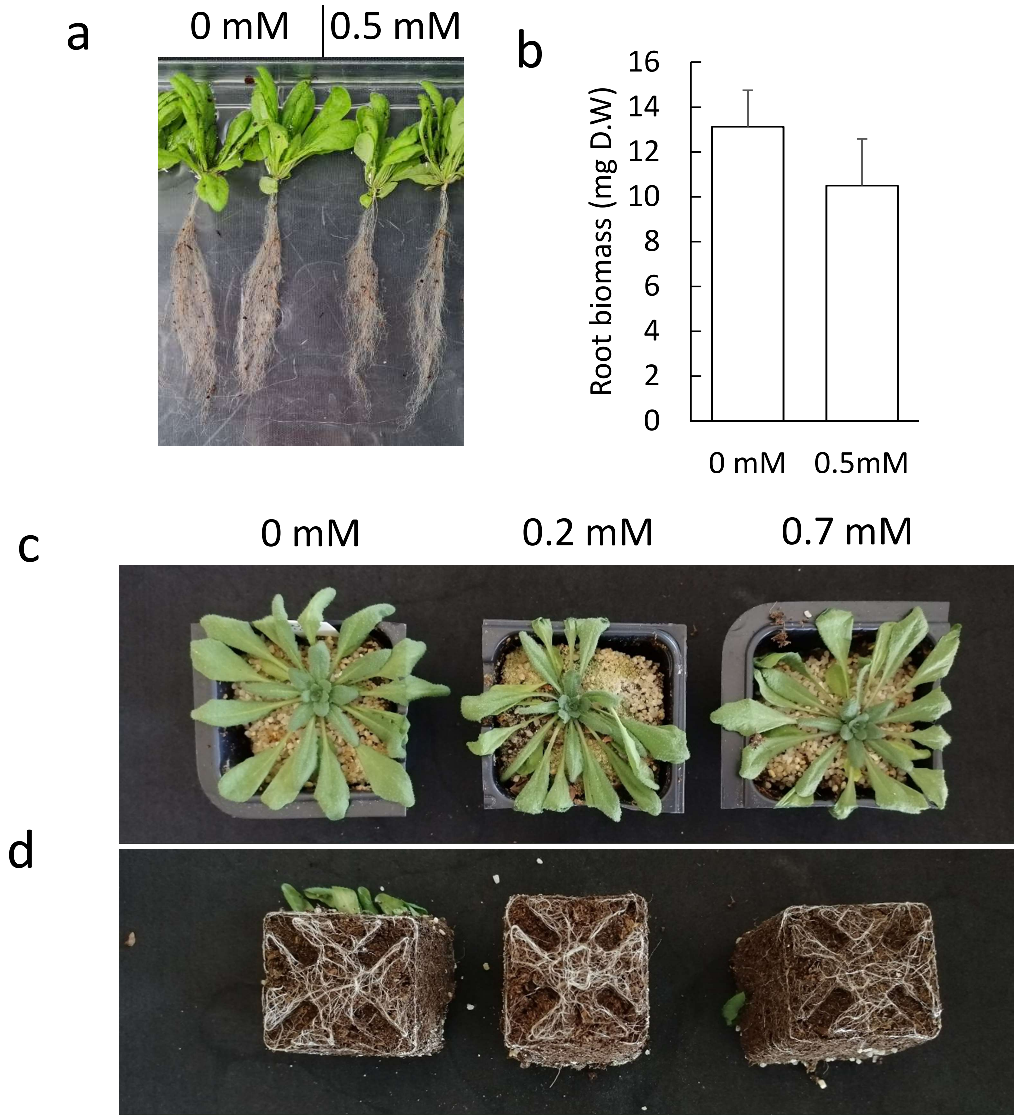


**Extended Data Fig. 7**. Spraying plants with β-CCA does not induce the same response as watering plants with β-CCA. a) Root growth of plants sprayed with 0 mM or 0.5 mM β-CCA. a) Root dry weight. c) Drought tolerance of plants sprayed with 0, 0.2 or 0.7 mM β-CCA. d) Picture of the roots at the bottom of the pots.

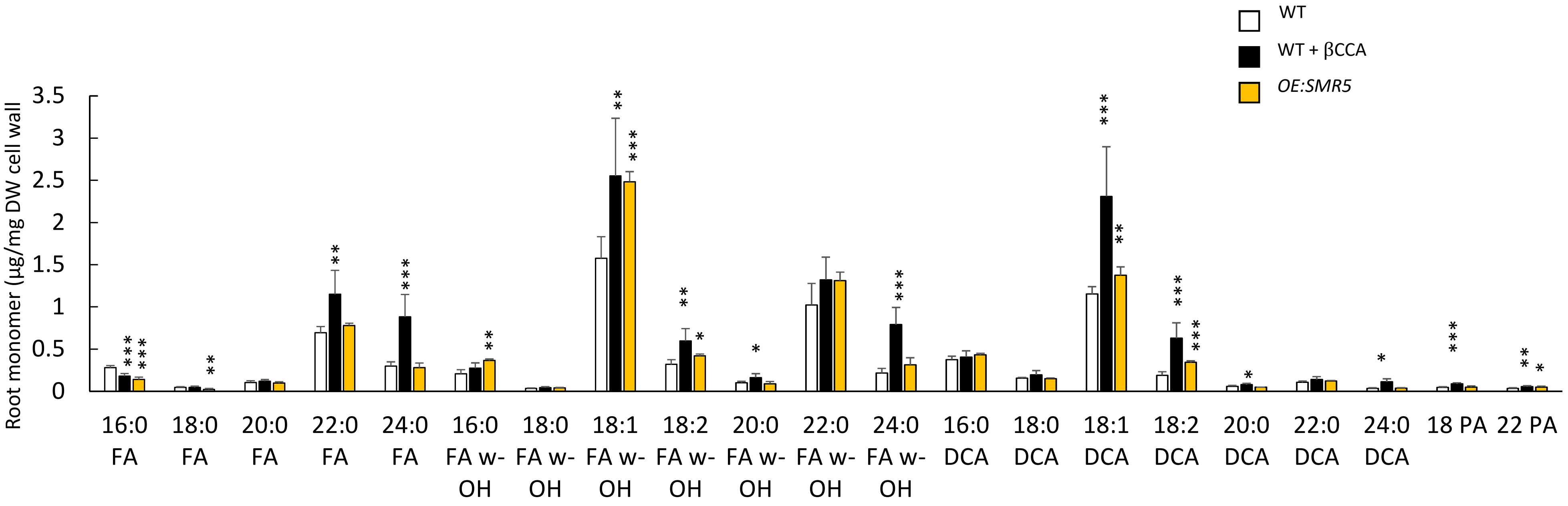


**Extended Data Fig. 8. Profile of root suberin monomers in WT, 75 µM β-CCA-treated WT and *OE:SMR5* seedlings**. Plants were grown in vitro of Agar solid medium. The values are means of 3 to 6 biological replicates. *, **, ***, different from WT at $P < 0.05$, 0.01 and 0.001, respectively (Student's t-test). FA, fatty acids; DCA, α,ω-dicarboxylic acids,; FA ω-OH, ω-hydroxy fatty acids; PA, primary alcohols.

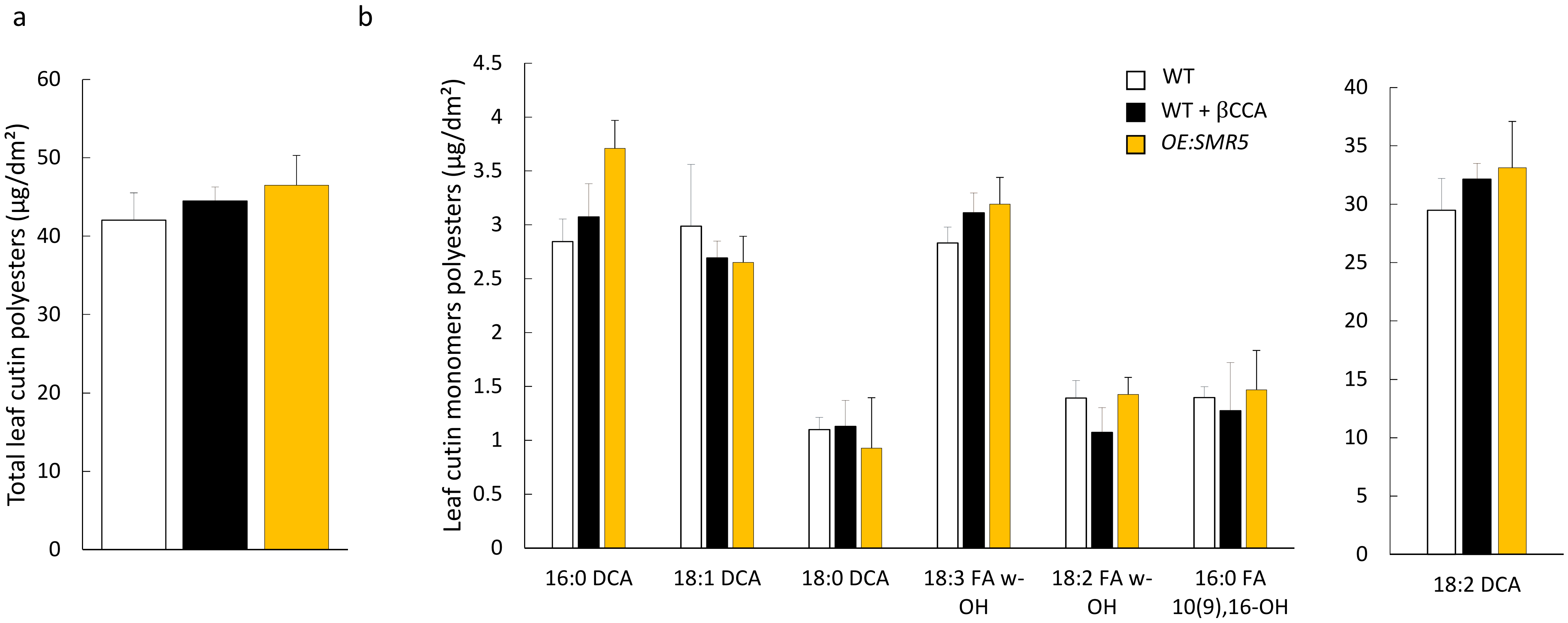


**Extended Data Fig. 9.** Leaf cuticle in Arabidopsis plants treated with 1.5 mM β-CCA or overexpressing *SMR5*. a) Total leaf cutin polyesters is calculated as the sum of each monomer shown in b). DCA, α,ω-dicarboxylic acids,; FA ω-OH, ω-hydroxy fatty acids.

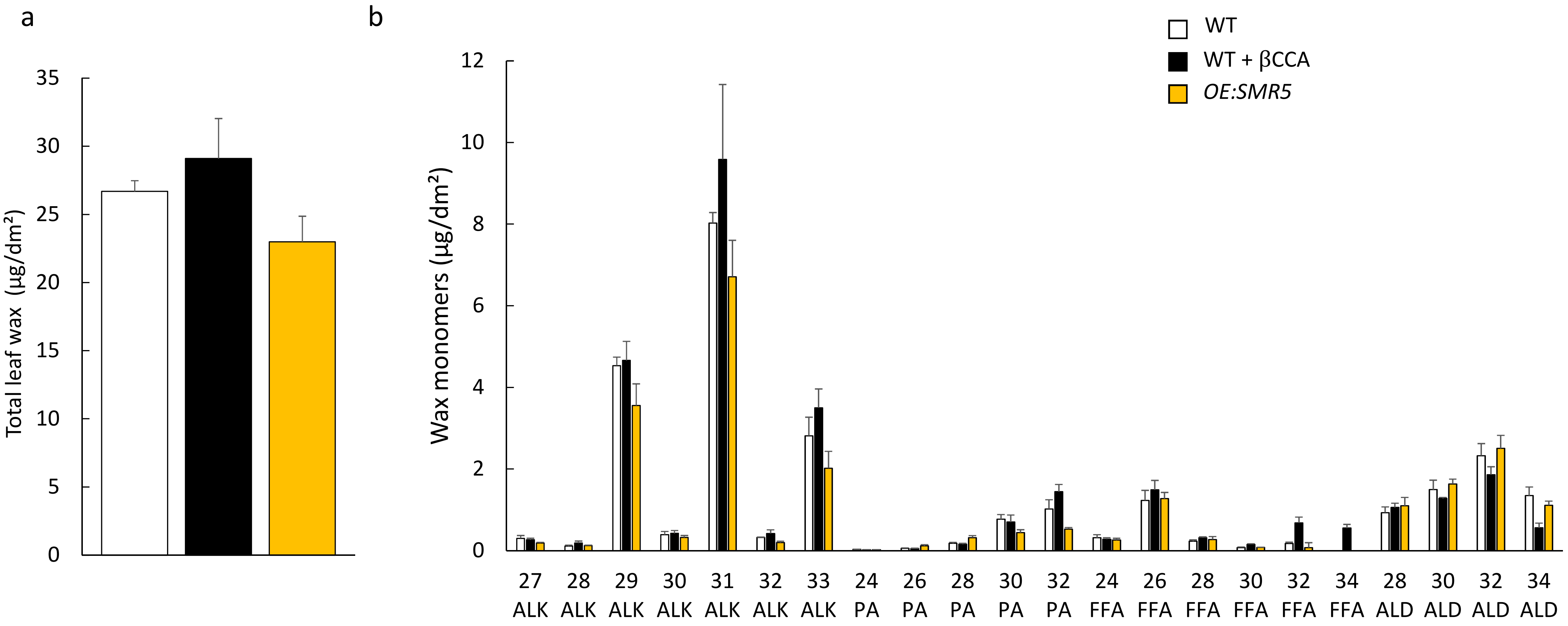


**Extended Data Fig. 10**. Leaf waxes in Arabidopsis plants treated with 1.5 mM β-CCA or overexpressing *SMR5*. a) Total leaf wax amount is calculated as the sum of each monomer shown in b). ALK, n-alkanes; PA, primary alcohols; FFA, free fatty aids; ALD, aldehydes.

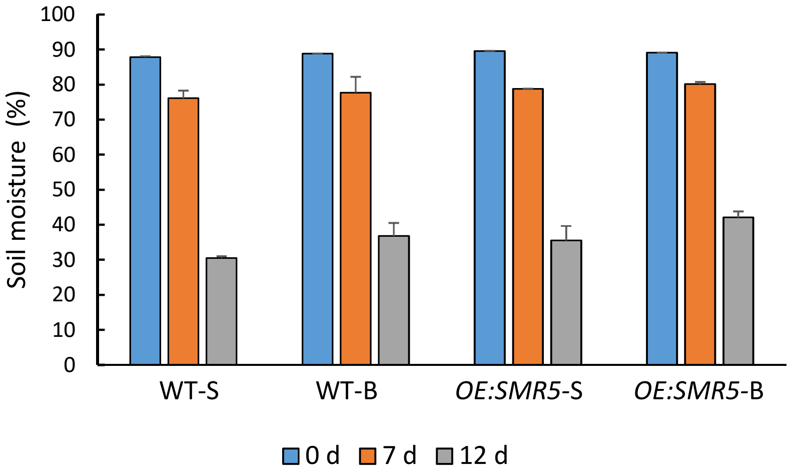


**Extended Data Fig. 11**. Soil moisture at the top (WT-S and *OE:SMR5*-S) and the bottom of the pots (WT-B and *OE:SMR5*-B) during water stress. WT and *OE:SMR5* plants were submitted to water deprivation for 0, 7 and 12 d.

**Supplemental Table 1.** List of primers used for qRT-PCR

| | | |
|---|---|---|
| **AT5G02220** | SMR4 | For: GAGGAAGACGGAGATGGCGG<br>Rev: AAAGTACCCGTTCCTCGGCG |
| **AT1G07500** | SMR5 | For: ACGACGGAGATACGGTGACG<br>Rev: CTCACCGGAGGTGGACAAGG |
| **AT3G27630** | SMR7 | For: AGCCGGTGAAGACGAAACTC<br>Rev: CGCCGTGGGAGTGATACAAA |
| **AT3G53090** | UPL7 | For: CTTCTGGGAGGTCATGAAAGG<br>Rev: CTCCAATAGCAGCCCAAAGAG |
| **AT4G26410** | UCP | For: CAGTTCCGCTCTATACAGGATTTC<br>Rev: GCGACACCAATCCCAATAGC |
